# Spike sorting biases and information loss in a detailed cortical model

**DOI:** 10.1101/2024.12.04.626805

**Authors:** Steeve Laquitaine, Milo Imbeni, Joseph Tharayil, James B. Isbister, Michael W. Reimann

## Abstract

Sorting action potentials (spikes) from extracellular recordings of large groups of connected neurons is essential to understanding brain function. Simulations with known spike times have driven significant advances in spike sorting, but present models do not account for neuronal heterogeneity and its effect on sorting accuracy. Here, we used a large-scale detailed cortical microcircuit model to simulate recordings, evaluate modern spike sorters, and link their performance to neuronal heterogeneity. We also exposed the network to various stimuli to investigate how sorting errors affect stimulus discrimination. Spike sorters successfully isolated about 10% of neurons within 50 µm of the electrode shank. This undersampling had no impact on stimulus discrimination ability. However, sorting biases related to firing rate, spike extent, synaptic type, and layer reduced its discrimination ability by nearly half. These findings show realistic models are a complementary method to evaluate and improve spike sorting and, hence, improve our understanding of neural activity.

## 1 Introduction

Neurons convey information by emitting sequences of electric signals called spikes. Emergent neural functions stem from sequences of spikes generated by large groups of interconnected neurons, called neuronal ensembles [1, 2]. These spikes can be recorded extracellularly, with electrodes positioned in the space surrounding the neurons. Recent advances in large-scale extracellular recordings with dense electrodes make it possible to capture the activity of neuronal ensembles with fine spatiotemporal resolution and scale. This provides a unique opportunity to explore the principles governing information processing by neuronal ensembles. In the last three decades, the field has advanced from tetrodes which captured only a few dozen neurons [3] to Neuropixels with 384 electrodes [4–6], increasing both the size of the recorded region and the density of recording sites.

At a fixed position, each neuron’s spikes have a characteristic, unique amplitude and shape. Spike sorting uses these unique signatures to detect spikes and assign them to source neurons. In large scale recordings, signals from multiple neurons close to the electrodes frequently overlap, a problem called spike collision. When many neurons are in the recorded field, as with modern, dense electrode arrays, this makes spike sorting both theoretically and computationally challenging. The rapidly increasing volume of data has compelled researchers to develop improved, automated spike-sorting algorithms, achieving state-of-the-art spike detection and neuron isolation with the Kilosort suite of computational algorithms [7]. To address the spike collision problem, Kilosort relies on template matching: detected spikes are iteratively subtracted from the raw signal after detection based on templates. This helps detect underlying spikes masked by the original spike. These combined hardware and software advances have elicited significant insights into how behavior is controlled by the activity of dynamic neural ensembles [2].

Studies using the Kilosort suite now typically report neuron yields of approximately 200 neurons for a Neuropixels 1.0 probe spanning the entire cortical thickness [1, 4, 8] in mice [4, 6], rats [8] and humans [9, 10] over periods of several days [11]. However, based on the density of cortical neurons and the rate of decay of spike amplitudes with distance to the electrodes one might expect 10 times more neurons to be isolated [1, 12]. For example, the most prominent neurons in the cortex, pyramidal cells, display sufficiently large spike amplitudes within 50 µm of an electrode site to be clearly detected, and so should in theory be easy to isolate using modern clustering methods [1]. It follows that spike sorting should accurately isolate about 800, 1300 and 1800 neurons in humans, rats, and mice respectively, given cortical thickness, neuron densities [13] and the geometry of a Neuropixels 1.0 probe [4]. The reported yields therefore correspond to only 25%, 16% and 11% of the theoretical maximum, highlighting substantial undersampling.

Applying spike sorting algorithms to small-scale simulations of extracellular recordings has led to substantial improvements in the algorithms, as this allows an objective comparison with the”ground truth” simulated spikes of the recorded neurons. These synthetic simulations have enabled an increase in the quality and number of isolated spike sorted units (i.e., the neuron yield). To improve the quality and yield of spike sorting algorithms, objective evaluations against “ground truth” have also been conducted using *paired extracellular and intracellular recordings* [8], where fractions of neurons not being isolated, i.e., error rates, as low as 5% have been reported for tetrodes [14]. However, this method is challenging for large dense electrode arrays, and may be limited to a small biased sample of neurons that exhibit the largest spike amplitudes [12]. Hybrid datasets combining spike sorted spike waveforms from ∼ 1000 neurons with simulated noise offer another option, again yielding an estimate of errors on the order of 5-10% [7, 15]. However, since the dataset is based on spikes detected from spike sorting and is manually curated [7], it may lack the full diversity and complexity of extracellular spikes *in vivo*, limiting the accuracy of the evaluation. In contrast, full synthetic simulations from a subset of 250 biophysically-detailed cortical neuron types, with independently generated spike trains [16, 17] do not rely on spikes obtained from spike sorting. Both hybrid synthetic and full synthetic simulations are flexible, allowing spiking features and noise levels to be easily tailored to test specific experimental hypotheses. However, although they allow tight control of relevant parameters, these models lack biophysical detail and do not capture the scale of real microcircuits, the diversity of spike waveforms arising from neuronal morphological heterogeneity [1] and the variability in spike firing patterns. Additionally, they do not incorporate synaptic connectivity, potentially failing to accurately replicate the statistics of temporally overlapping spikes [18]. These simplifications may inflate sorting accuracy and yield in synthetic extracellular recordings, which could explain the disparity between the experimental yield and the neuron yield predicted by the synthetic datasets.

More specifically, the discrepancy in neuron yield could be attributed to systematic biases in spike sorting algorithms. Indeed, the evaluation of spike sorters on synthetic ground truth datasets has revealed systematic biases, which can compromise the quality of isolated spike trains and how well they reflect true neural activity [16, 19]. For example, overmerging the spike trains of multiple neurons into a sorted unit or oversplitting a neuron’s spike train across multiple sorted units [19] can obscure the information content in both individual neuronal and collective population responses [20, 21]. Additionally, background noise may produce spurious sorted units [7, 16, 20]. These biases could be explained by the sensitivity of spike sorters to certain anatomical and physiological neuron features, which, if insufficiently characterized due to lack of a realistic ground truth, may reduce yield. Such biases may preferentially sample certain neuronal populations and overlook others, such as sparsely firing neurons [6, 21, 22]. They may also distort critical information about neural activity [23], and obscure the true nature of the neural code at both single neuron and population levels [20, 21, 23]. Furthermore, mean firing rates estimated from sorted spike trains are often used for the calibration of models of brain circuitry [24–27]. Biases may degrade these estimates, leading to less realistic models. Without realistic ground truth datasets to better characterize how spike sorters handle anatomical and physiological neuron features, these limitations remain particularly pronounced in large-scale, dense recordings where manual curation of millions of spikes from hundreds of neurons is impractical. A detailed network model with high biological validity would support automated curation aimed at minimizing spike sorting errors and enable comparisons between ground truth neuronal population representations and those derived from spike sorting.

Here, we tested whether modern spike sorting algorithms are biased towards certain anatomical and physiological neuron features in a biophysically-detailed model of an entire cortical column of the rat primary somatosensory cortex [28, 29], providing a new ground truth for evaluating and improving spike sorting algorithms. The biophysically-detailed model was constructed in a bottom-up manner, i.e., by carefully modeling and independently validating the individual components in separate, in-depth peer reviewed studies before combining them into a single model [30–36]. As such, extracellular activity emerges from biophysical principles, rather than being generated from a biased sample of waveforms. Neurons of diverse morphological and electrical classes were accurately distributed in accordance with anatomical laminar density profiles and modeled as spatially extended multi-compartmental models with their electrical models fit and validated against single cell protocols. The simulated extracellular activity emerges from a generative model that captures the full heterogeneity of cortical tissue. During calibration of background input and validation of emerging population activity, potential overestimates of mean firing rates from sorted spike trains were taken into account. This enabled layer-wise responses of excitatory and inhibitory populations to single whisker deflection stimuli to closely match *in vivo* references on a millisecond time scale. Furthermore, we validated that the model produced realistic *in vivo* extracellular recording traces, spikes, and sorted neuronal firing statistics.

We evaluated the yield, accuracy, and sensitivity to neuronal features of the complete suite of Kilosort algorithms: Kilosort (1, 2, 2.5, 3, 4) [7]. We also assessed herdingspikes, an alternative spike sorter also designed for large-scale multi-electrode arrays [37]. We found that the number of units sorted from the simulated data was comparable to yields reported *in vivo* for all tested spike sorters, but also that it represented only 15% of the theoretical yields, i.e., of neurons within 50 µm of the electrode shank. Most isolated neurons were either incomplete or merged with multiple units. Spike sorting accuracy was ten times lower in the biophysical models than in the less detailed full synthetic simulation [16], demonstrating a substantial effect of neuronal anatomical and physiological features on spike sorting. In a regression analysis, sorting accuracy was positively related to small electrode distance, high firing rate, large spike spatial extent, excitatory neuron type and being within cortical layer 5. There was a pronounced over-representation of layer 5 pyramidal cells compared to other layers across all spike sorters, while interneurons were occasionally over-represented with older spike sorting methods.

In addition to simulating rest activity, we simulated the response of primary somatosensory cortex to whisker deflection. We thereby demonstrated that some of the biases detailed above significantly affect the recovered representations of stimuli. Biased sampling reduced a linear classifier’s ability to distinguish between eight simulated stimulus inputs to the microcircuit based on population responses by nearly 50%, while spike assignment biases lowered performance by 30%. Surprisingly, the significant undersampling observed did not impact the circuit’s stimulus discrimination ability. Overall, these findings suggest that while spike sorting preserves a degree of stimulus information, it also produces an imperfect representation of neural activity. Based on the generated ground truth data, we further provide a model that automates the curation of high-quality single neurons based on their spike features, achieving nearly 90% precision and 60% recall. Our openly available dataset, models, and analyses provide a biophysically realistic foundation for improving and automating spike sorting as datasets continue to expand.

## 2 Results

### 2.1 Model simulation

To address the lack of biological detail of current synthetic simulations [7, 16], we used a detailed and heterogeneous model of cortical tissue incorporating diverse neuronal morphologies, firing patterns, and synaptic connections [28, 29]. The model extends the work of [30] to an atlas-based model of the full non-barrel primary somatosensory cortex. We simulated a 400 µm diameter sub-volume comprising 30,190 neurons distributed over all six cortical layers. Neurons were connected by a total of 36.7 million synaptic connections, with each neuron associated with a reconstructed biophysical morphology. The model captures a significant degree of biological variability with 60 morphological types (18 excitatory and 42 inhibitory, Fig. 1a), 11 firing classes (Fig. 1b), and 30,190 neurons with unique reconstructed morphologies and morpho-electric combinations (Supplementary Fig. 1a).

**Fig. 1.**
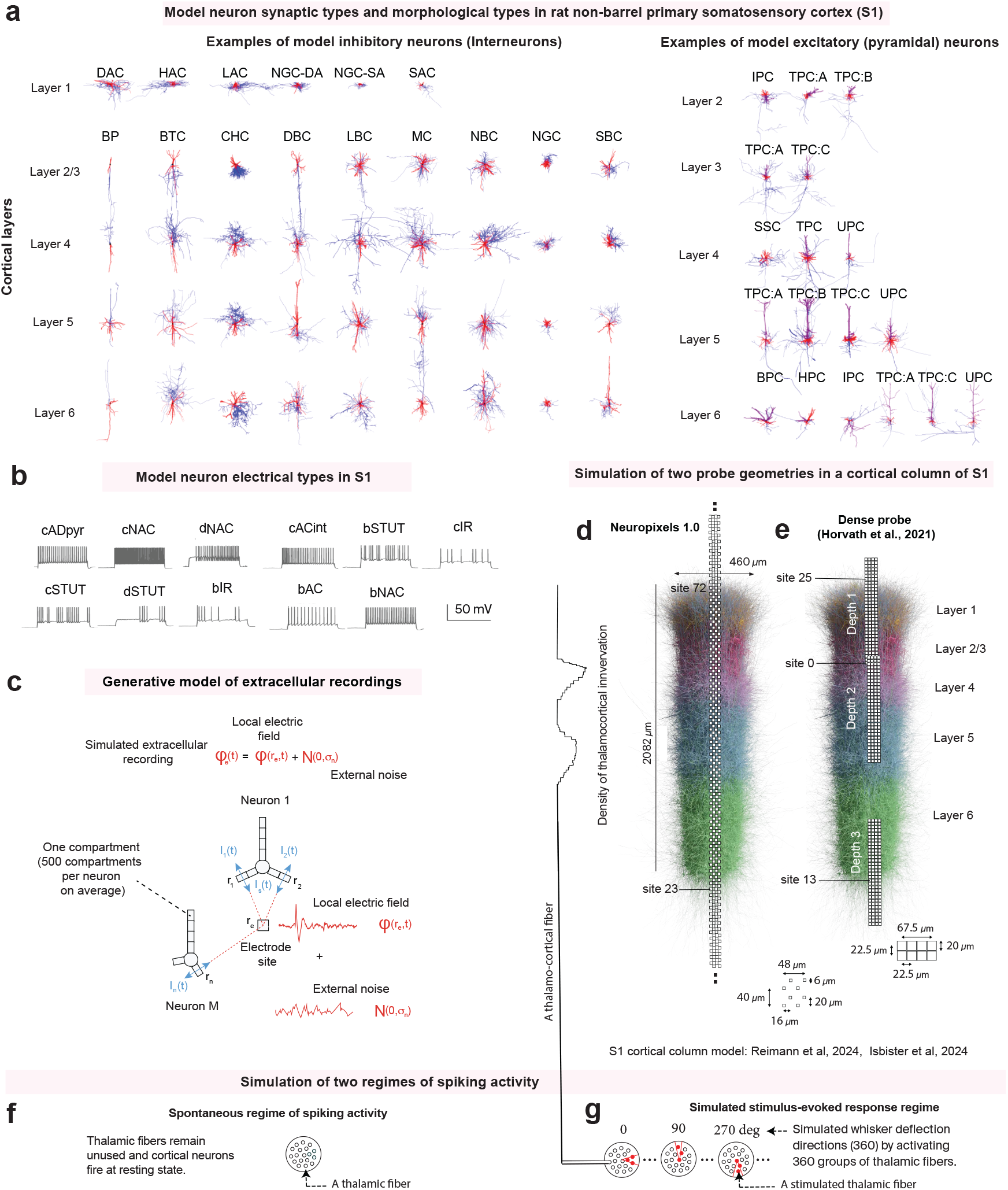
Biophysical models and experiments. **a**. Example of modeled inhibitory (left) and excitatory neuron morphologies (right) in per cortical layer [28]. Axon in blue, dendrites in red. **b**. 11 different firing patterns, **c**. Generative model of the extracellular recordings [39], **d**. Bouton distribution across cortical depth (line ascending from panel **g** [28, 30, 48]) assuming one afferent fiber from ventral posteromedial (VPM) thalamic nucleus, overlaid on randomly chosen neurons. Simulated **d**) Neuropixels 1.0 and **e**) Horvath probe [41]. Electrode widths but not inter-electrode distances, are multiplied by 3 compared to the column’s scale. In bottom panels, distances are multiplied by 3. **f**. Spontaneous activity regime in which thalamic fibers remained unused and cortical neurons fired at resting state, **g**. The experiment was used to validate the model against recordings of anesthetized rats [8, 41]; Evoked regime in which 360 groups of fibers were activated to simulate neural responses to whisker deflections in 360 directions repeated 50 times for 200 ms. This experiment was designed to assess the effect of spike sorting on stimulus cortical representation. Morphologies: DAC (HAC, LAC or SAC): Descending (Horizontal or Large or Small) Axon Cell, NGC-DA (or NGC-SA): Neurogliaform Cell with dense (or slender) axonal arborization, BP: Bipolar Cell, BTC: Bitufted Cell, ChC: Chandelier Cell, DBC: Double Bouquet Cell, LBC (NBC or SBC): Large (Nest or Small) Basket Cell, MC: Martinotti Cell, NGC: Neurogliaform Cell, IPC: Inverted Pyramidal Cell, TPC (or UTPC): Tufted (or Untufted) Pyramidal Cell, TPC:A (or TPC:C): Large (or Small) Tufted Pyramidal Cell, TPC:B: Early Bifurcating Pyramidal Cell, SSC: Spiny Stellate Cell, BPC: Bipolar Pyramidal Cell, HPC: Horizontal Cell, see [28].

To simulate realistic electrical signals in the extra-cellular space surrounding neurons, we used a well established approach, which divides neurons into smaller sections (∼ 500 compartments per neuron on average) and calculating the contribution of each section to the electric signals in extracellular space [38] (Fig. 1c). This allowed us to describe in detail how different parts of the neuron – like the cell body (somata), the branches (dendrites), and the starting point of the axon – shape the electric field, that is, the electrical signals in extracellular space. In these detailed simulations, we calculated the extracellular electrical signals by modeling the dendrites and the axon initial segment as lines emitting currents into the extracellular space (Fig. 1c, line-source equation [39, 40]), and the soma as a single point (see point-source equation in Methods). This generative model allowed us to simulate extracellular recordings at the sites of electrodes on different electrode probe designs used in real-world recordings (*in vivo*).

A recent study found that improving the spatial resolution of recorded electric signals around neurons can enhance spike sorting in mice by detecting small neurons with small electric fields (small spatial footprints) [6]. To explore the effect of electrode density on spike sorting accuracy, we tested two types of recording devices (probes): one with wide coverage and electrodes closely packed together, the Neuropixels 1.0 probe [4] (Fig. 1d), and another with more closely spaced electrodes but a smaller overall area, the dense probe introduced in [41] (Fig. 1e). In this study, we did not use the probe configuration as in [6], with both full coverage of the cortical thickness and high electrode density because of its high computational requirement. The Neuropixels 1.0 probe was made of a single shank with 384 square electrodes arranged in 4 staggered columns and distanced at 26µm (bottom, Fig. 1d), as described in [4]. For this probe, we simulated recordings and compared them against Neuropixels data from the anesthetized rat from [8]. The denser probe (bottom, Fig. 1e) had more restricted coverage of the cortical column. For this probe, we simulated recordings at three depths (top, Fig. 1e), replicating the setup used in the anesthetized rat study by [41], reconstructing their electrode locations as closely as possible (Supplementary Fig. 1b). The probe had 128 electrodes arranged in 4 columns, horizontally and vertically separated by 22.5 µm. To our knowledge, these were the only two publicly available dense recording datasets from the rat primary somatosensory cortex. Using identical probe configurations as *in vivo* enabled us to directly compare/validate the model’s extracellular activity against experimental data and assess how spike sorting performance depends on electrode layout and spatial resolution. However, it should be noted that we did not attempt to simulate the exact experimental conditions of those references (such as anesthesia), hence differences in the dynamic states were expected.

We simulated both spontaneous and stimulus-evoked neural activity (Fig. 1f) to assess the robustness and accuracy of spike sorting across different activity regimes and to evaluate the effect of information loss on stimulus recovery using spike sorting algorithms (Fig. 1g). The model included thalamic input fibers, which synapse onto cortical neurons with layer-wise innervation profiles and were used here to apply whisker-like stimuli by sending spike trains down a subset of fibres (Fig. 1g-d). In the spontaneous condition, the thalamic input fibers remained unused and neurons fired at resting firing rates. In the evoked condition, stimuli were repeatedly delivered through the thalamic input fibers to mimic higher and more structured, activity-driven firing. We used 360 distinct *stimulus patterns*, each associated with a subset of the thalamic fibers. For each stimulus run, thalamic fibers were activated for 200 ms. To provide sufficient data, 50 stimulus runs were presented to the model for each of the 360 stimuli.

To evaluate whether using a more detailed and heterogeneous representation of cortical tissue had an impact on spike sorting performance, we compared sorting accuracies and yields between the biophysical simulations and the simpler full synthetic ground truth simulation described in [16], referred to here as the “synthetic model”.

### 2.2 Model validation

Spike sorters’ ability to detect and isolate a neuron’s spike depends on several features of the recorded trace. It increases when the spike amplitude is large relative to other spikes of smaller amplitude produced by neurons at about 150 µm (multi-unit activity) from the electrode [1]. It also increases when spike amplitude is large relative to the amplitude of the background electrical signal produced by neurons located beyond 200 µm (the background noise) [1]; i.e. when the spike has high signal-to-noise ratio. We thus assessed the validity of the Neuropixels and dense recordings simulations by comparing these features of the generated traces, the shape of its extracellular spikes, and the firing statistics of the sorted spikes to those described in the Marques-Smith Neuropixels data [8] and the Horvath dense probe data [41] respectively. However, our goal was not to precisely replicate these studies, but rather to ensure that our simulated extracellular traces accurately reflected the key characteristics of spike amplitude, spike shape, background noise, and multi-unit activity within the physiological range observed *in vivo*.

#### 2.2.1 Voltage trace components

As *in vivo*, the preprocessed traces contained spikes with high signal-to-noise ratio, multi-unit activity, and background noise (Fig. 2a-f). Single-unit waveforms were apparent at nearby sites (Fig. 2g-l) and exhibited correlated spatiotemporal dynamics similar to *in vivo*. As expected, the denser probe (Fig. 2k, l) sampled the electric field with higher spatial resolution than the Neuropixels probe (Fig. 2g-j).

**Fig. 2.**
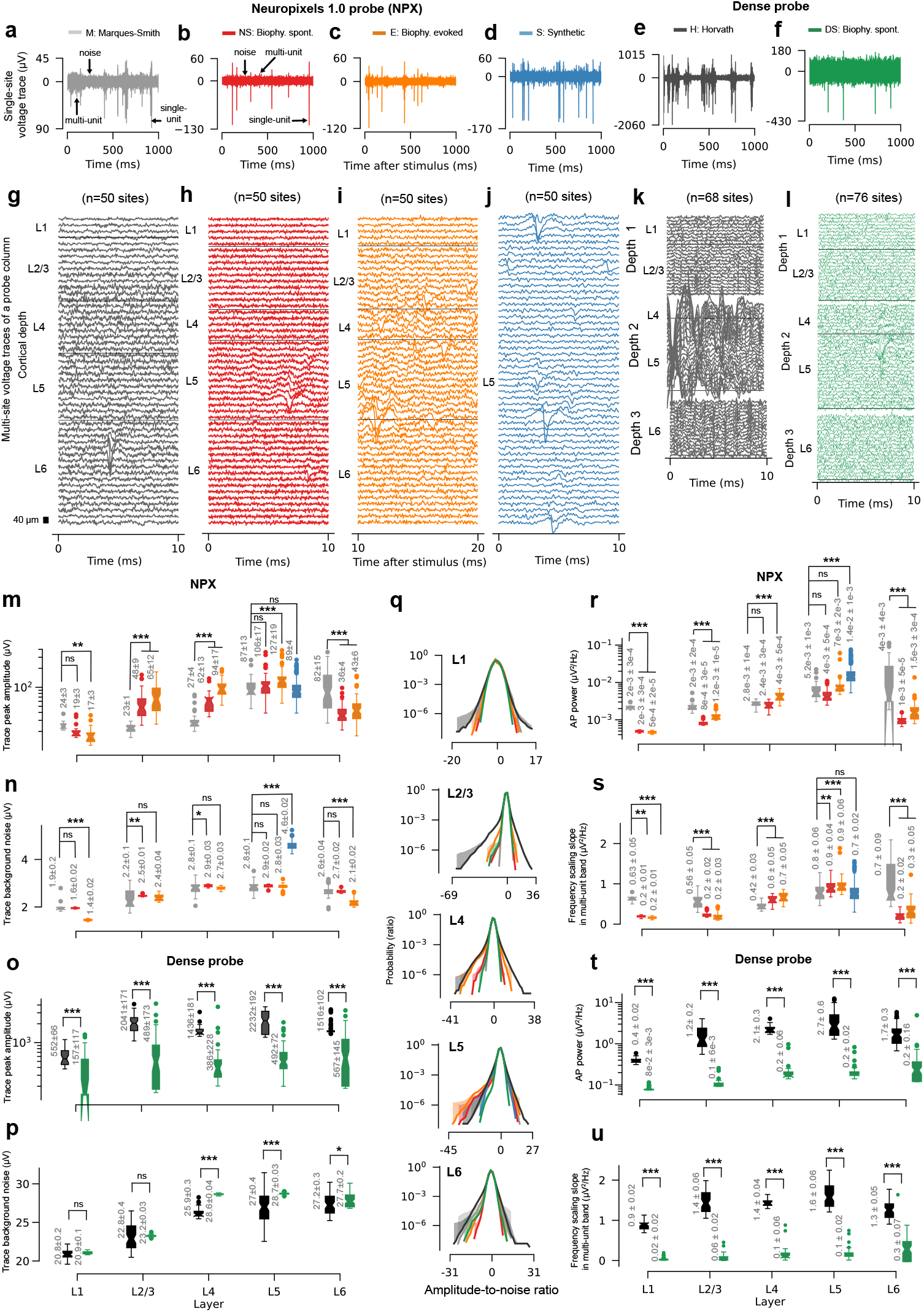
Voltage trace validation. **a-f**. Single-electrode traces in layer 5 for **a**) Neuropixels 1.0 recordings in anesthetized rats [8], **b**) the biophysical model of spontaneous recordings and **c**) 10 ms after simulated whisker deflections on the same site, **d**) from a synthetic simulation [16], **e**) from anesthetized rat recordings with the denser probe [41] (insertion depth 2, Fig. 1e) and **f**) for the biophysical simulation of the denser probe spontaneous recordings. **g-l**. Multi-site traces, color-matched to **a-f**, shown for one probe column (50 among 96 sites shown for Neuropixels). Peak amplitude distributions for **m**) Neuropixels and **o**) denser probes. Background noise distributions (noise is lowest median absolute deviation (MAD) over one-second bins) for **n**) Neuropixels and **p**) denser probe. **q**. Median distribution of the trace amplitude-to-noise ratios across sites and upward 95% confidence interval for 100 bins. Amplitudes were divided by the MAD of each site. **r**. Boxplots of AP power and **s** fitted frequency scaling slope (*α*) from the power law 1*/f* ^*α*^ [43] for the Neuropixels probe for frequencies between 0.3-2 kHz (single spike) and between 0.3-6 kHz (multi-unit spiking) respectively. **t-u**. Same as **r-s** for the denser probe, at depth 1 for L1 and 2/3, at depth 2, for L4 and 5, and depth 3 for L6. Measurements in **m-u** are across electrodes from the entire recordings. p-values: ns, *p >*= 0.05, *, *p <* 0.05, ***, *p <* 0.01, ***, *p <* 0.001 from Kruskal-Wallis test with Dunn’s multiple comparison and Holm-Sidak adjustments.

#### 2.2.2 Voltage trace amplitude-to-noise ratio

Spikes with low signal-to-noise ratio (SNR) are challenging for spike sorters to detect and isolate, as their amplitude and shape are often obscured by background noise. To enhance the fidelity of the biophysical models, a model of external sources of noise was fitted layer by layer, and the trace extrema were adjusted to match *in vivo* (see Methods). We then assessed whether their SNR distributions aligned with *in vivo* ranges and reflected layer-specific variations across cortical tissue. Spike SNR is typically defined as the peak amplitude of the sorted spike divided by the amplitude of the background noise. To obtain a measure of SNR that is not biased by spike sorting, we compared the distribution of peak amplitudes, background noise levels and amplitude-to-noise ratios (ANR) of the electrode traces. The ANR is calculated by dividing the trace amplitudes by the median absolute deviation (MAD).

The median peak amplitudes and background noise levels of the biophysical traces tended to differ from Marques-Smith and Horvath data (χ^2^ between 13 and 138, most *p* < 0.05, df = 2, kruskal-Wallis test with Dunn’s multiple comparison test between experiments, Fig. 2m-p). Precisely, the median peak amplitude of the neuropixels biophysical simulations was 34% higher *in vivo*, on average across layers and experiments (40% across layers 1 and 6 and 113% across L2/3 to 5, most *p* < 0.01, Fig. 2m). Their overall median background noise level was also lower than *in vivo*, but only by 1.5% (9% lower in layers 1 and 6 and 3% higher in layers 2/3 to 5, most *p* < 0.05, Fig. 2n). For the dense probe, simulated median peak amplitude was 73% lower than *in vivo*, on average across layers and experiments (all *p* < 0.001, Fig. 2o) and simulated median noise was marginally higher than *in vivo*, on average by 1.8% across layers (most *p* < 0.05, Fig. 2p). Nonetheless, the three models reproduced the increase in median peak amplitudes and background noise levels observed *in vivo* across layers (χ^2^ between 34 and 232, *p* < 0.001, df = 4, Kruskal-Wallis between layers) and both metrics were generally within *in vivo* range.

The distributions of ANRs observed *in vivo* and in the models showed a wide dynamic range, with alignment of data points between -10 and 10 times the noise level (Fig. 2q), particularly in layers 1, 5, and 6. Despite deviations from *in vivo* due to difference in peak amplitudes, simulated ANRs remained within the *in vivo* range. These findings demonstrate that the biophysical models effectively capture key aspects of *in vivo* trace signal-to-noise ratio, including their dynamic range and layer-specific trends, despite differences in peak amplitudes.

#### 2.2.3 Spiking components

A high level of spiking activity should increase the temporal overlap of spikes, making it more difficult for spike sorting algorithms to distinguish and isolate individual spikes. To ensure that the spiking activity in our simulations remained within the *in vivo* range, we used power and frequency scaling as indirect measures of spiking activity. These measures do not depend on spike sorting, thus providing unbiased estimates of the theoretical number of spikes present in the data. We decomposed the voltage trace into its constituent frequencies and compared the amount of frequencies (power) associated with single-spike waveforms (AP power [42], within 0.3-2 kHz) in the simulations to Marques-Smith and Horvath data. If all spikes had the same shape and amplitude, and did not overlap throughout the recording, AP power would be directly proportional to the number of spikes [42]. Power typically decays with increasing frequency according to a power law 1/f^α^ [43] where α quantifies the decay rate. This decay rate reflects the relative contributions of single-spike waveforms and multi-unit spiking frequencies to the electrode trace in the spiking frequency band (0.3-6 kHz). After confirming that the electrode traces’ power spectral density were well fitted by power laws in the spiking frequency band (Supplementary Fig. 2a), we compared simulated and *in vivo* slopes α within this frequency range.

The median AP powers of the biophysical traces and the fitted power law slopes tended to differ from *in vivo* (χ^2^ between 25 and 167, most *p* < 0.01, df = 2, kruskal-Wallis test with Dunn’s test between experiments, Fig. 2r-u). The biophysical models reproduced *in vivo*’s median AP power for the Neuropixels probe better than it did for the dense probe. The median AP power of neuropixels biophysical traces was 54% lower than *in vivo*, on average across layers and experiments (69% lower across layers 1, 2/3 and 6 and 12% higher across layers 4 and 5, most *p* < 0.001, Fig. 2r). Median fitted power law slope was also lower than *in vivo* by 59%, on average across layers and experiments (70% lower across layers 1, 2/3 and 6 and 31% higher across layers 4 and 5, all *p* < 0.01, Fig. 2s). The dense probe simulation consistently displayed much weaker power than *in vivo* across layers compared to the neuropixels biophysical simulations, on average 90% lower than *in vivo* (all *p* < 0.001, Fig. 2t). Median power law slopes were also consistently weaker than *in vivo* across layers compared to the neuropixels biophysical simulations (on average 93% lower than *in vivo*, all *p* < 0.001, Fig. 2u). However, the three models successfully captured the increase in AP power and power law slope across layers observed *in vivo* (χ^2^ between 56 and 144, most *p* < 0.001, df = 4, Kruskal-Wallis between layers) and both metrics were generally within *in vivo* range. Overall, these results highlight the biophysical models’ ability to also reproduce features of *in vivo* spiking activity, capturing the layer-specific variations of single-spike and multi-unit spiking activity, despite some discrepancies in power and slope magnitudes.

We note that in Horvath data but not in Marques-Smith’s, the traces were dominated by high-amplitude slow (0.5 - 1.5 Hz) and δ waves (1 - 4 Hz) characteristic of slow-wave sleep [44] and visible to the naked eye across layers (Supplementary Fig. 2b and e). The biophysical models did not replicate these oscillations (Supplementary Fig. 2c, d, f) under the current parametrization but are able to do so with alternative meta-parameter combinations (results not shown). We assumed that these differences had a negligible effect on spike sorting, as we high-pass filtered the traces above 300 Hz as is typically done to eliminate slow oscillations prior to spike sorting.

#### 2.2.4 Sorted unit spike shapes

Crucially, spike waveform variations - such as reduced amplitude [1], broader spikes [45], and increased distortion with distance from the soma [1, 45] - can significantly affect the ability of sorting algorithms to detect spikes. If these sources of variability are not adequately represented in synthetic models, they may contribute to the yield discrepancy and systematic biases observed in spike sorting. Past experimental studies have identified five spike temporal profiles on single electrodes with shapes that are consistent across species (in humans, see panel Fig. 3a copied from [9], and in cats, see panel Fig. 3b copied from [46]). Although the most recently released Kilosort4 sorter has been shown to outperform all other sorters [7], we adopted the same approach as in [9] for direct comparison: both the simulated data and the Marques-Smith data were sorted with Kilosort3, and spike waveforms were extracted using spike-triggered averaging. The simulated temporal waveforms qualitatively matched the five shapes reported in cats [46] and humans [9] (Fig. 3c vs. a-b) and their variation with the location of the electrode aligned with previous simulations [45]. Spike waveform variation exhibited three features with distance to the somata: redistribution from negative to positive components, decreased peak-to-peak amplitude, and widening of spike amplitudes (Fig. 3d). The peak-to-peak amplitude of the spikes rapidly decayed between nearby electrodes, dropping by more than half within 75 µm (Fig. 3e), therefore making the summed extracellular potentials difficult to isolate at 75 µm from the somata. The extracellular trace was strongly distorted by noise beyond 200 µm (Fig. 3f) in agreement with [1].

**Fig. 3.**
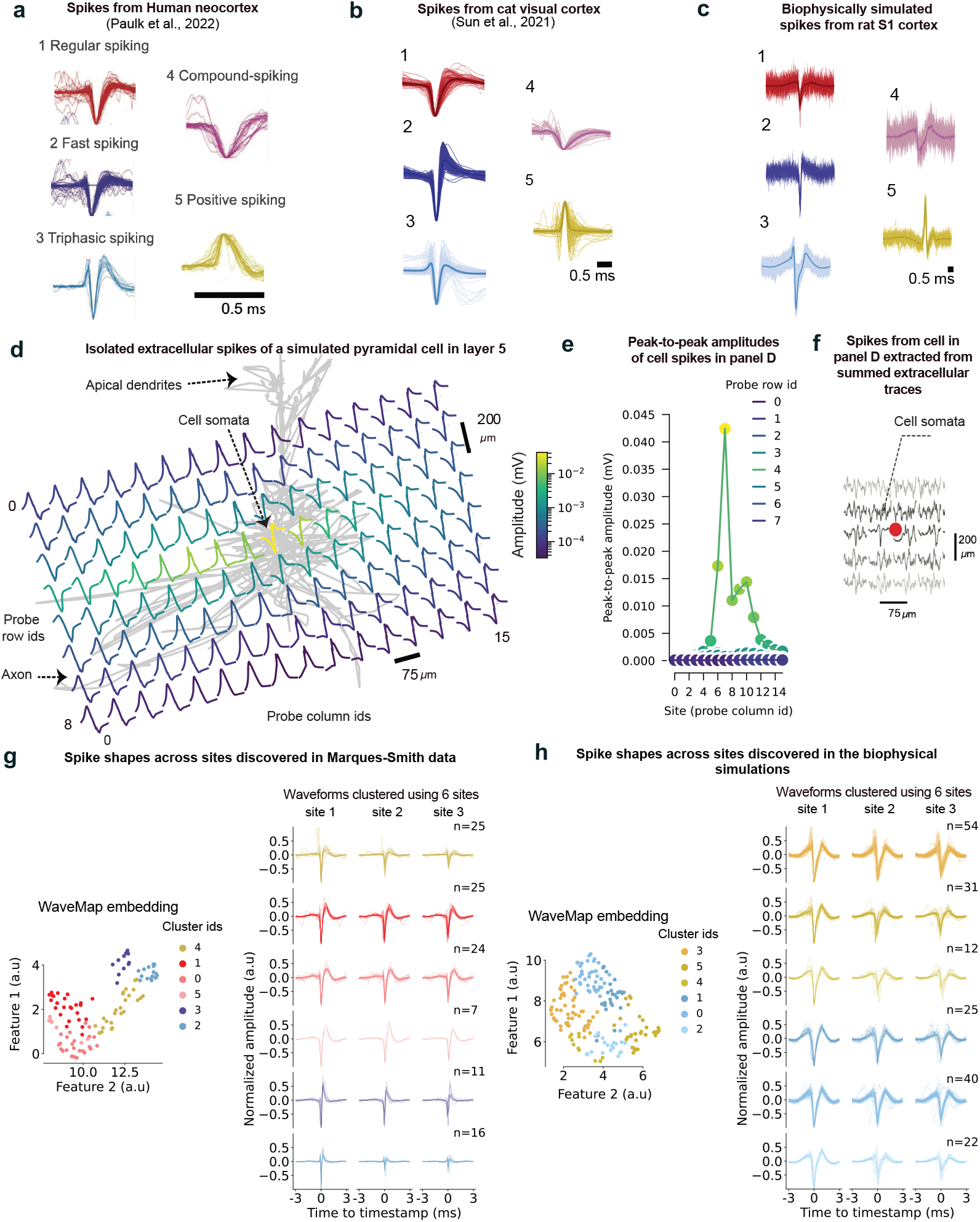
Validation of spike spatiotemporal profiles. **a**. Five types of spike temporal profiles sorted with Kilosort3 from Neuropixels 1.0 single-electrodes in the Human neocortex (panel adapted from [49]), **b**. Five similar types found in the cat visual cortex (panel adapted from [46]) and sorted with Kilosort3 in the **c**) biophysically simulated Neuropixels recording in the spontaneous regime. **d**. Isolated extracellular recording of the most active biophysically simulated pyramidal cell in layer 5 at 128 electrode locations organized according to a grid layout. The electrodes and spikes are spatially aligned with the cell morphology (grey) **e**. Peak-to-peak voltage amplitudes of cell spikes from **d** recorded at sites in row 4 and at all column locations. The peak-to-peak amplitude reaches its maximum at column site 8, near the cell’s somata and rapidly drops at the two nearest recording sites, 75 µm away from the somata. **f**. Extracellular recording of the cell in **d** including the contribution of all the microcircuit cells and from the 20 sites nearest to the somata. The cell’s spike is severely distorted by noise at 200 µm from the somata. **g**. Embeddings in two dimensions of sorted single-unit spike waveforms obtained from *Wavemap* [47]. To enable comparison with [49], spikes were sorted with Kilosort3 from Marques-Smith Neuropixels recordings [8] and from **h**) the neuropixels spontaneous biophysical simulation. The average spike waveform of each neuron (lines) corresponding to each identified cluster (colors) are shown in **g** and **h** for the three nearest electrodes with the largest average spike amplitude. n indicates the number of neurons per cluster. *Wavemap* identified six spike shape clusters that are qualitatively similar between the *in vivo* recordings and the simulation.

We further validated that the biophysical models not only replicated the general features of spike waveform variability but also captured the diversity of spatiotemporal profiles observed in *in vivo* data. We used *WaveMap*, a new machine learning tool designed to group sorted spikes based on the shapes of their waveforms across different electrodes [9, 47]. We then compared the spike profiles identified to those discovered in the Marques-Smith dataset (Fig. 3g). Consistent with the experimental data, the algorithm identified six distinct profiles from the simulations (Fig. 3g, h). These profiles were visible on the isolated extracellular traces of both model excitatory (Supplementary Fig. 3a) and inhibitory neurons (Supplementary Fig. 3b), and the shape of the waveforms varied with the locations of the electrodes.

#### 2.2.5 Sorted unit firing rates

Firing rate is a key factor influencing spike sorting accuracy [7, 16]. To assess its variation across cortical layers and how well simulations replicate *in vivo* firing rate distributions, we used Kilosort4, the most advanced spike sorter [7] to ensure the comparisons reflected the highest achievable match. To improve unit isolation, Kilosort4 leverages the fact that neurons have a refractory period — a brief interval (1–5 ms [7]) following a spike during which they cannot fire again. It differentiates ‘single-units’ from ‘multi-units’ by classifying sorted units with minimal violations of the neuronal refractory period as ‘single-units’ (a “contamination rate” less than 0.1, see Method). We compared the firing rate distributions of ‘single-units’ across three datasets: Marques-Smith Neuropixels recordings [8], biophysical simulations of Neuropixels recordings, and Buccino’s full synthetic simulations of Neuropixels recordings [16] (Fig. 4). We excluded comparisons involving the denser probe simulations with *in vivo* data because their lower peak amplitudes (Fig. 2o), lower levels (Fig. 2u) and different compositions (Fig. 2v) of spiking activity, although within the *in vivo* range, could skew sorting yield and quality. We analyzed the first ten minutes of each recording, which was the longest common recording duration. Both simulated and *in vivo* data showed lower single-unit yields in superficial layers, with significantly higher yields deeper layers, prompting a focus on layer 5 and 6. *In vivo*, the yields in these layers were at least 20 times that in layer 4, and no units were found in layers 1 and 2/3, a trend also reflected in the simulated data (Fig. 4a, see unit count).

**Fig. 4.**
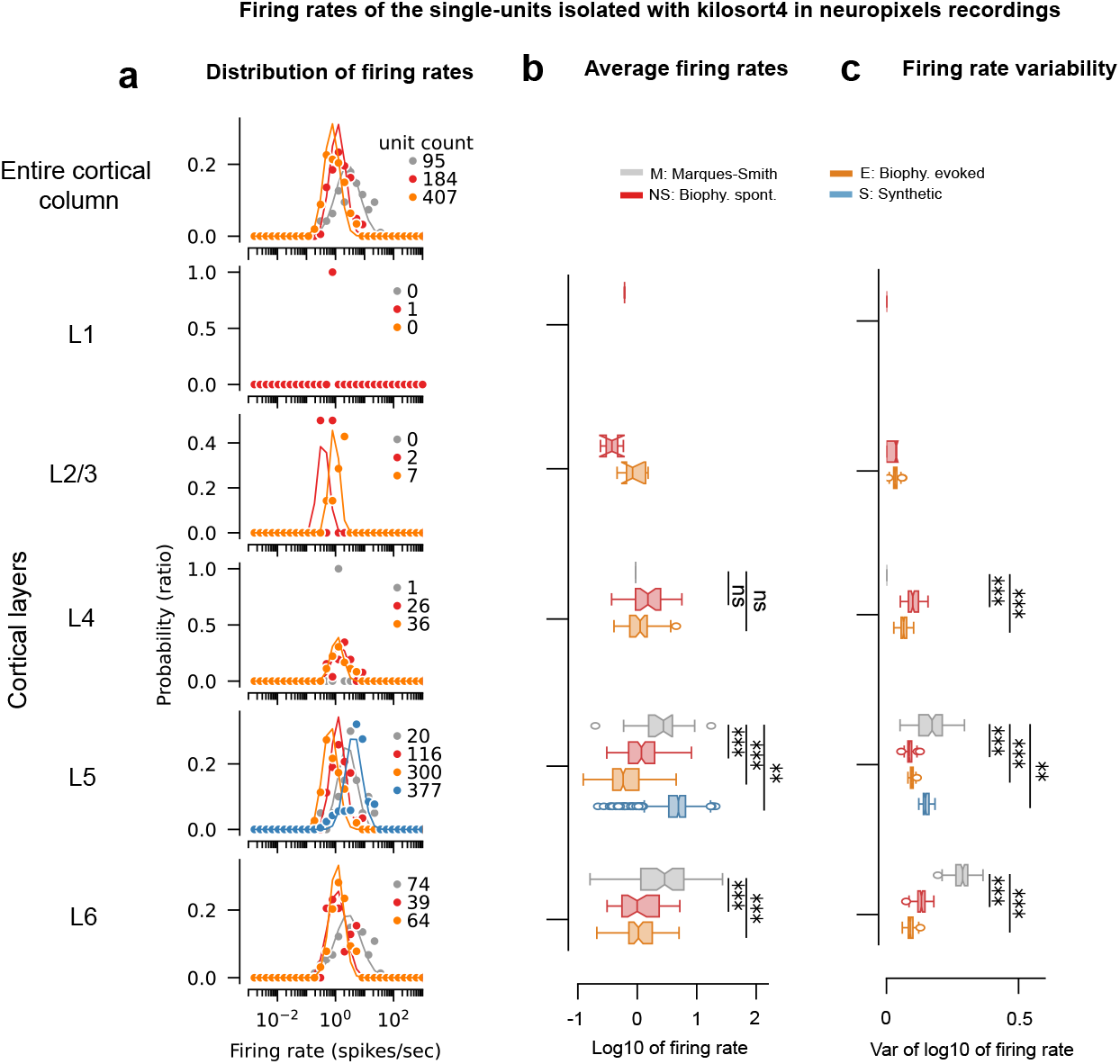
Validation of single-unit firing rate distributions sorted with Kilosort4 in simulated Neuropixels recordings. **a**. Distribution of the firing rates of the single-units sorted with Kilosort4 in Marques-Smith data (grey dots), the spontaneous (red dots) and evoked (orange dots) biophysical simulations. Distributions are also reported for Buccino’s synthetic model of recordings in layer 5 (blue dots). The distributions were fitted with lognormal distributions (color-matched lines), which provided a good qualitative match to the data. **b**. Boxplots of log-transformed firing rates. Firing rates were log-transformed with the decimal logarithm to facilitate the visualization of lognormally distributed data. **c**. Boxplots of the variances of distributions of the firing rates generated from 100 bootstraps of the decimal logarithm of the firing rates. The boxplot notches represent the 95% confidence intervals of the median and variances respectively. Statistical significance of the null hypothesis that the simulated median firing rates and variances are the same as *in vivo* data are reported with ns, *,**,***, which indicate *p >*= 0.05, *p <* 0.05, *p <* 0.01, *p <* 0.001 using the Mann-Whitney U test.

The biophysical simulations successfully captured the major trends in firing rate statistics, despite minor differences. Like Marques-Smith’s sorted unit firing rates, the firing rates from the spontaneous (red dots, Fig. 4a) and evoked (orange dots) biophysical simulations were distributed log-normally across layers (red and orange lines). The overall median firing rate averaged over layer 5 and 6 differed but only by a small margin (Marques-Smith: 2.8 ± 1% (95% CI) spikes/secs, spontaneous biophysical simulation: 1.2 ± 0.2%, evoked simulation: 0.7 ± 0.1% and Buccino’s full synthetic simulation: 5 ± 0.4% (95% CI)). Specifically, the biophysical simulations had lower median firing rates (Fig. 4b, *p* < 0.001, Mann-Whitney) and distributions were narrower (Fig. 4c, *p* < 0.001) than *in vivo* across layers. The synthetic model distribution also had a lower variance in layer 5 (all *p* < 0.01) but it displayed a significantly higher median firing rate than *in vivo* (p < 0.01, Mann-Whitney).

Older spike sorters also tended to produce firing rates than were more homogeneous than *in vivo*. While newer sorters like Kilosort3 produced log-normal distributions (Supplementary Fig. 4a) with median firing rates similar to *in vivo* in the spontaneous regime (Supplementary Fig. 4b, *p* > 0.15, Mann-Whitney), older sorters often generated bimodal distributions with peaks aligned between *in vivo* and all three simulations. Across both spontaneous and evoked conditions, median firing rates showed no clear trend across older sorters (Supplementary Fig. 4b), while variances in simulations tended to be reduced (Supplementary Fig. 4c, *p* < 0.01), highlighting a consistent but marginal gap in capturing the full range of neural activity seen in *in vivo* recordings.

Overall, these shared trends demonstrate that the biophysical simulations closely capture key aspects of *in vivo* variability, despite the absence of biological inputs from non-simulated regions that may account for a lower overall activity level and reduced variability.

### 2.3 Model predicts significant ‘single-unit’ undersampling and assignment biases

High-density probes typically allow the isolation of approximately 200 neurons per probe [1, 4, 6, 8–10]. After validating the biophysical simulations, we tested whether spike sorting produced comparable yields to those observed *in vivo*. To enable a fair comparison with the *in vivo* yield, we discarded signals from electrodes located in the white matter: counting only ‘single-units’ isolated by Kilosort4 on electrodes in the cortex in Horvath and Marques-Smith data. Having access to ground truth data also permits evaluating the quality of the isolated units, to characterize systematic spike sorting biases and to improve the isolation of high-quality units.

The spontaneous biophysical simulations produced a yield of ‘single-units’ that more closely matched *in vivo* data compared to the synthetic model, despite producing more sorted ‘single-units’ than *in vivo*. We found that for spontaneous activity recorded with Neuropixels and sorted by Kilosort4, on average 41% of sorted units were ‘single-units’. This proportion differed between experiments (Fig. 5a, χ^2^(4, *N* = 5, 241) = 43.9, *p* < 0.001). At least twice more ‘single-units’ were detected from the simulations than *in vivo*, with the largest difference being 206 vs. 93 for the dense probe (Fig. 5a, 182 and 193 vs. 96 for Neuropixels). Yet the fractions of ‘single-units’ were similar (χ^2^(2, *N* = 1, 212) = 2.0, *p* = 0.37 for Neuropixels, χ^2^(1, *N* = 920) = 1.8, *p* = 0.18 for the dense probe). In contrast with the *in vivo* yield, the majority of units sorted from the synthetic traces were ‘single-units’, four times more than *in vivo* (S: 377 vs. 163, χ^2^(1, *N* = 740) = 4.0, *p* = 0.042). The greater similarity of the biophysical model to *in vivo* yield could be attributed to differences in spike timing statistics. In the synthetic model, spike timing was generated with an independent Poisson process while they result from synaptic connectivity in the biophysical models, which will produce more spike collisions.

**Fig. 5.**
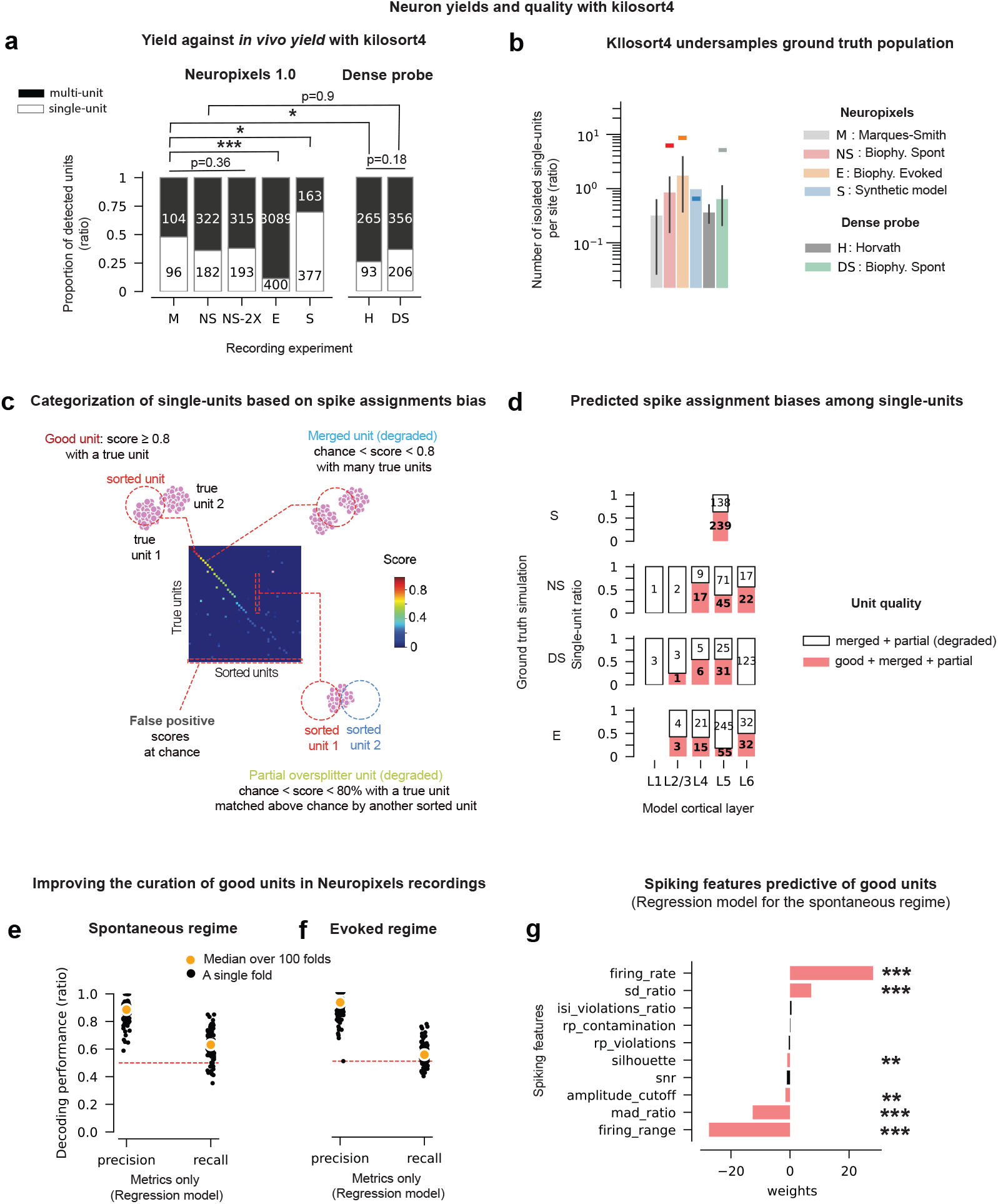
Model predicts undersampling and spike assignment biases by Kilosort4. **a**. ‘Single-units’ vs. ‘multi-units’ ratio from Marques-Smith data (M), Horvath data (H), spontaneous (NS) and evoked (E) Neuropixels biophysical simulations, Buccino’s simulation [16] (S), and dense probe biophysical simulation (DS). Spike waveform shortening in NS (NS-2X) to match *in vivo* duration as done in [7]. *: *p <* 0.05, **: *p <* 0.01, ***: *p <* 0.001, Pearson’s *χ*^2^ goodness-of-fit test, **b**. ‘Single-unit’ yield per electrode. Yield per electrode’s medians (bars) and 95% CI (error bars) across layers; median theoretical yields per site (colored horizontal lines, NS: 6.9, NE: 8.4, S: 0.65, DS: 5.3), i.e., active ground truth neuron count within 50 µm divided by site count and averaged across layers. **c**. ‘Single-unit’ categorization based on the type of spike assignment biases. ‘Good’ units display matching scores with ground truth neurons above 80%; ‘Merged’ units have scores below 80% but above chance with multiple ground truth neurons; ‘Partial’ units have scores between chance and 80% with a ground truth neuron also matched above chance by another unit; False positives have a score at chance (see method). **d**. Ratios of spike assignment biases (colors) by layer (x-axis) and simulation (row panels). Numbers are counts. A ‘good + merged + partial’ unit (pink) displays above 80% sorting accuracy with a ground truth neuron; it oversplits that neuron, which is matched by another sorted unit with above chance accuracy, and it matches another neuron with above chance accuracy. **e-f**. Precision and recall performance of a lasso-regularized logistic regression model that classifies ‘good’ vs. degraded units using spiking quality metrics in **e**) NS and **f**) NE. Cross-validation done with 100 bootstraps of 75%train - 25% test samples; Performance for 25% of test units (black dots) and their median (orange dots). **g**. Model weights for NS (see quality metrics description in Method). Weights significantly different from 0 (pink); quality metrics were z-scored to compare their magnitudes. *: *p <* 0.05, **: *p <* 0.01, ***: *p <* 0.001, Student’s t-test.

The model predicts that the number of sorted units should increase at least three times in a stimulus-evoked regime. Although the ratio should strongly shift to ‘multi-units’, a key prediction is that the single-unit yield should double (E:400 vs NS:182, Fig. 5a).

To understand whether higher-resolution sampling of the electric field influences the isolation of ‘single-units’, we compared the number and types of units detected by Kilosort4 for the two simulated probe designs. Kilosort4 detected more units from the dense probe than the Neuropixels probe (H:358 vs M:200 units and DS:562 vs. NS:504 units respectively, Fig. 5a). These were mostly ‘multi-units’, which were twice more frequent for the dense probe *in vivo* (M:104, H:265, χ^2^(1, *N* = 558) = 6.6, *p* = 0.01). The model did not reproduce this effect (NS:322, DS:356, χ^2^(1, *N* = 1066) = 0.004, *p* = 0.9).

Despite a higher-resolution sampling of the electric field, the dense probe did not produce a higher single-unit yield than Neuropixels, which cannot be attributed to a lower amplitude-to-noise (ANR) ratio since the *in vivo* dense recording had the highest ANR (Horvath, Fig. 2e, q). These findings suggest that while denser probe geometries detect more units, the additional captured units may be difficult to isolate and instead resemble ‘multi-units.’

A recent study suggested that the extended duration of biophysically simulated extracellular spikes, compared to *in vivo* spikes, reduces the quality of spike sorting by increasing false positives [7]. To mitigate this, the authors proposed downsampling the simulated traces by a factor of two. We implemented this approach and, in agreement with their findings, observed a 6% increase in single-unit yield (experiment NS-2X in Fig. 5a). However, the modest magnitude of this improvement suggests that while spike duration may influence sorting accuracy, it is not the primary limiting factor in our simulations. Instead, probe geometry and activity regime played a more significant role.

The observed single-unit yields are significantly lower than the number of neurons with sufficiently large spikes within 50 µm of the probes [1, 12]. We quantified the gap between the observed and the theoretical yield. For the biophysical models, we counted the number of units within 50 µm of the probes that fired at least once (theoretical yield). To enable comparison between the two probes, which have different numbers of electrodes per layer, we report the median yield per electrode averaged over layers. For Buccino’s model the theoretical yield was simply 0.65 units per site (250 model units and 384 electrodes). For the biophysical models (red and orange bars), the raw single-unit yields were an order of magnitude lower than the theoretical yields (color-matched horizontal bars, Fig. 5b) and were not significantly different between probes or spike sorters (Supplementary Fig. 5a). In Buccino’s full synthetic model (blue bar, Fig. 5b), Kilosort4 produced more ‘single-units’ than the proportion of modeled neurons per site, indicating the production of redundant units. These observations were true for most tested sorters (Supplementary Fig. 5a), except for Kilosort2, which detected a large number of units [16]. The original Kilosort also apparently achieved maximal yield but because it does not curate ‘single-units’ from ‘multi-units’, it is possible that it also has low single-unit yield.

Since the simulator provided the actual spike times for each simulated neuron, we were able to assess the quality of the isolated ‘single-units’ and identify significant issues with their accuracy. We matched isolated ‘single-units’ to corresponding *true units* by finding the simulated neuron that maximizes the agreement score of their spike trains [16] (for details, see methods). The agreement score is a similarity metric for pairs of spike trains that reach a value of 1.0 for a perfect match. How close the maximizing neuron gets to an agreement score of 1.0 can also serve as a measure of sorting quality for a given sorted unit. This also allowed us to classify sorted units based on the types of errors that occurred during sorting (Fig. 5c): If, for a single spike sorted unit for example, the second-best matching ground truth neuron has an agreement score close to the best ground truth neuron, it is likely that the unit is a ‘merged unit’ (blue in c), i.e., representing the union of the spike trains of both neurons. The majority of the isolated ‘single-units’ were less than 80% accurate and degraded across layers (Fig. 5d, white area) and spike sorters (Supplementary Fig. 5b). They contained spikes from multiple true units (‘merged unit’) and missed more than 20% of the matched true units’ spikes (‘partial or oversplitter unit’ in green, Fig. 5c). Kilosort4 and 3 did not produce false positives (black area, Fig. 5c,d and Supplementary Fig. 5b) but older kilosort versions did, in agreement with previous reports [7, 16].

### 2.4 Improving ‘single-unit’ quality

To improve quality, the output of spike sorters is typically manually curated. To automate this arbitrary [50] and laborious process for Kilosort4, we developed a Lasso-regularized logistic regression model to classify ‘good’ and degraded ‘single-units’ based on ten spiking features. These features or quality metrics [20] were selected for their comprehensive statistical descriptiveness of a unit’s spike waveforms and firing properties [16], computational efficiency, and applicability to most units (182/184 in the spontaneous Neuropixels simulation).

We introduce a new metric, the spike mad_ratio (mean absolute deviation ratio), which quantifies the contamination of single units by other units. Defined as the ratio of the mean absolute deviation of a unit’s spike trough (negative peak) to that of the surrounding background noise, this metric exhibited the strongest correlation with sorting accuracy (R = − 0.46, Pearson correlation), outperforming all other quality metrics in predicting sorting performance.

To maximize model performance and generalization while ensuring simplicity and interpretability, we applied Lasso (Least Absolute Shrinkage and Selection Operator) regularization. This approach mitigates overfitting by adding an L1 penalty to the logistic regression loss function, which shrinks feature coefficients and sets some to exactly zero. By selecting only the most relevant spiking features, Lasso not only enhances model accuracy and robustness but also improves interpretability by reducing complexity to the essential predictors.

We assessed model performance through cross-validation, training on 75% of the dataset and testing on the remaining 25%. Given the imbalanced proportions of ‘good’ and degraded units, we evaluated precision (the proportion of predicted ‘good’ units that were actually ‘good’) and recall (the proportion of actual ‘good’ units correctly identified).

The model demonstrated very strong and robust performance, correctly classifying ‘good’ single-units in both spontaneous and evoked regimes. It achieved precision scores of 89% ± 2% (95% CI) and 93% ± 2%, and recall rates of 61% ± 3% and 55% ± 2%, in the spontaneous (Fig. 5e) and evoked simulations (Fig. 5f), respectively. While the recall was slightly lower, with the model missing less than half of the high-quality units, the high precision indicates that Kilosort4’s single-unit features still carry valuable information for improved curation.

Furthermore, the model identified firing rate statistics (firing rate and range, *p* < 0.001, Student’s t-test) and the spike mad_ratio (p < 0.001) as the strongest predictors of high-quality units, with these three features influencing sorting accuracy by an order of magnitude more than others (Fig. 5g).

We evaluated whether incorporating the full waveform, alongside the ten quality metrics, could improve curation performance. Given the high-dimensional nature of spike waveforms—240 timepoints over a 6 ms period sampled at 40 kHz—one might expect that using all available spike information would enhance both precision and recall. To reduce waveform dimensionality for each single-unit, we employed CEBRA, a neural network autoencoder with strong generalization capabilities that outperforms other dimensionality reduction methods without making assumptions about data distribution [51]. CEBRA also allows for the use of supervised labels to enforce strong separation between labels in the low-dimensional embeddings. We utilized this feature to enhance the distinction between ‘good’ and degraded unit spikes. We then trained a k-Nearest Neighbor classifier to classify units based on the embeddings produced by CEBRA. Results are reported for three-dimensional embeddings as performance did not improve for higher dimensions.

While incorporating the full waveforms did improve recall (82% ± 3% and 90% ± 2% for the spontaneous (Supplementary Fig. 5c) and evoked (Supplementary Fig. 5d) regimes, respectively), it also led to a significant reduction in precision (61% ± 3% and 55% ± 3%), resulting in many more false positives. We ensured that CEBRA’s loss function always converged so this lower precision was not due to insufficient training (Supplementary Fig. 5e). Rather, it likely stems from the model overfitting noise introduced by the random component of the full waveforms.

Given the computational cost of processing the waveforms and training and cross-validating the CEBRA model, we conclude that the logistic regression approach, which relies solely on quality metrics, is more efficient and provides better precision for curation.

### 2.5 Biological factors of sorting failures

The biophysical models successfully replicate several characteristics of high-density extracellular recordings. The biological validity of the models allowed us to further explore how factors such as spike sorting algorithms, probe design, and the anatomical and physiological properties of model neurons impact spike sorting accuracy and contribute to ‘single-unit’ undersampling and spike assignment biases. We defined the sorting accuracy of a ground truth neuron’s as its agreement score (proportion of correctly detected spikes) with its best matching sorted ‘single-unit’. Ground truth spikes were considered detected if they were matched by a sorted unit’s spikes within a 1.3 ms time window (Δ_*t*_ = 1.3 ms, see Method). This window was chosen to maximize sorting accuracy for Kilosort4 (Supplementary Fig. 6a) while avoiding inflated scores due to random matches (Supplementary Fig. 6b).

As reported in previous studies [16], most spike sorters detected a high proportion of ground truth neurons with accuracies above 80% (ranging from 50% of neurons for Herdingspikes to 99% for Kilosort2) in the full synthetic simulation (Fig. 6a). Similar high sorting accuracies are reported in hybrid synthetic datasets [7]. In contrast, the biophysical simulations, which modeled the anatomical and physiological heterogeneity of the cortical tissue, produced notably lower proportions of well-sorted ground truth neurons (Fig. 6a). On average, only 15% of the neurons within 50 µm of the electrodes were correctly sorted across the simulations (Fig. 6b, χ^2^(5, *N* = 3, 190) = 4.9, *p* = 0.43). This discrepancy highlights the impact of modeling tissue heterogeneity in more realistic simulations.

**Fig. 6.**
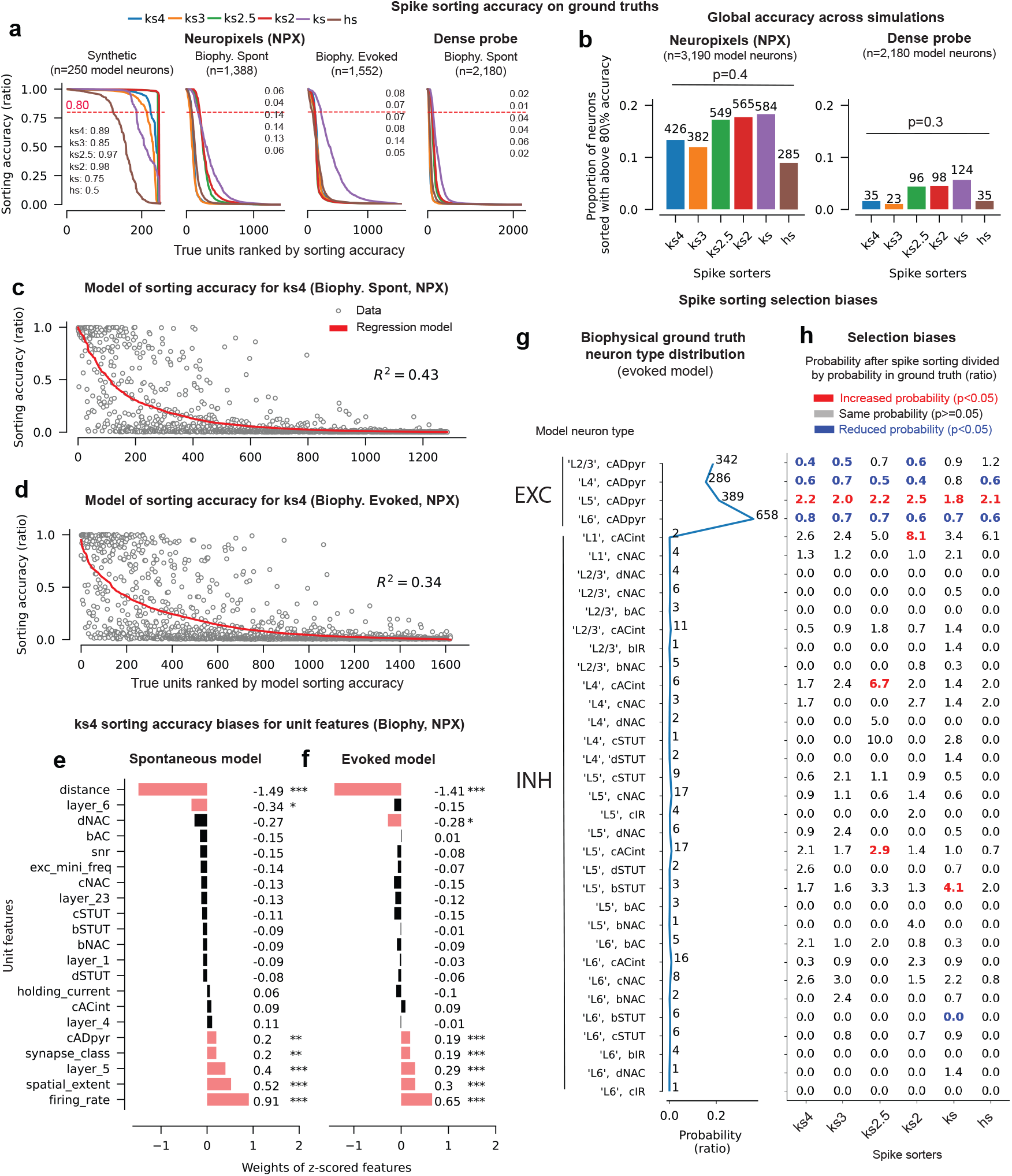
Sorting accuracy and sensitivity to neuron anatomical and physiological features. **a**. Ground truth neuron sorting accuracy, i.e., the proportion of ground truth neurons (see annotations) sorted with above 80% accuracy within 50 µm of the electrodes, **b**. The total sorting accuracy summed across datasets, i.e., the number of well-detected neurons across ground truths divided by the number of neurons within 50 µm of the electrodes across simulations. For the dense probe, numbers were pooled across depths. P-values from *χ*^2^ goodness-of-fit tests. Biophysical ground truth neurons’ sorting accuracies against best-fitted regularized fractional regression models (red line) for Neuropixels simulations of the **c**) spontaneous and **d**) evoked regimes, sorted with Kilosort4. *R*^2^ are models’ explained variance, **e**,**f**. Model weights. Weights in pink are significantly different from 0 and numbers indicate weight amplitudes. *: *p <* 0.05, **: *p <* 0.01, ***: *p <* 0.001. **g**. Probability distribution of neuron types defined by their synaptic class, layer and electrical type, in the evoked biophysical ground truth. Number annotations indicates the count of each model neuron type **h**. Ratios of the probability of occurrence of model neuron types in the sorted data to their probability of occurrence in the ground truth. Each sorted single-unit was first assigned the biophysical features of its best ground truth match and we calculated the ratio of ground truth neurons’ probability of occurrence after sorting and in the ground truth. Fisher’s exact test, two-tailed. The 10-minute evoked simulation at 40 kHz was used in panels **a** and **b**. To link neuron selection biases to stimulus representation quality (Fig. 7), the 1-hour evoked simulation at 20 kHz was analyzed in panels **d, f**, and **g-h**.

Despite its ability to sample the electric field with higher spatial resolution, the dense probe achieved a ‘single-unit’ yield comparable to that of the Neuropixels probe (Fig. 5a-b) and the quality of these single-units was surprisingly lower. To account for the fact that the dense probe did not span the entire cortical column like the Neuropixels probe, leaving portions of layers 5 and 6 unsampled (Fig. 1e), we normalized the number of well-sorted ground truth neurons by the number of neurons within 50 µm of the electrodes, eliminating potential layer sampling biases. Even after normalization, the dense probe yielded five times fewer neurons sorted with above 80% accuracy across spike sorters compared to the Neuropixels probe (on average 3% vs. 15%). Together with Fig. 5a, this result suggests that while kilosort4 captures more spiking activity from denser electrode geometries - leading to an increase in detected multi-units - it struggles to effectively isolate high-quality single-units from the increased activity. Given its higher sorting accuracy and full cortical column coverage, we focused our analyses on the Neuropixels probe.

We leveraged the detailed morphological, anatomical, and physiological descriptions of our ground truth model neurons to systematically characterize how neuron features influence sorting accuracy. To achieve this, we used a lasso-regularized fractional logistic regression model, which is well-suited for predicting continuous proportions (see Methods). As in our curation model, lasso regularization was used to enhance generalization and interpretability. However, unlike the curation model, this analysis included 21 ground truth neuron features—which are not accessible to experimentalists—to directly assess their impact on sorting accuracy.

The model provided a good fit to the data (*R*^2^ = 0.42 and *R*^2^ = 0.34 in the spontaneous (Fig. 6c) and evoked (Fig. 6d) regimes respectively, Mcfadden pseudo *R*^2^), identifying seven significantly predictive features out of 21 in both the spontaneous and evoked conditions. The dominant factor was distance to the nearest electrode, which exhibited an absolute coefficient 1.6 times larger than the next most influential feature (firing rate) and 25 times larger than the weakest predictor (’dSTUT’-firing type). This finding aligns with expectations: neurons closer to an electrode should be sorted with higher accuracy (Fig. 6e,f)).

In agreement with prior synthetic and hybrid simulations, the model also predicts that sorting accuracy improves with higher firing rates ([7, 16]) and with greater spike spatial extent across electrodes—consistent with past simulations [7] and *in vivo* studies ([6]). However, our model quantifies novel biases in their relative importance: distance to the electrode influences sorting accuracy three times more than firing rate, and firing rate is twice as influential as spike spatial extent.

One might expect interneurons to be sorted with greater accuracy than pyramidal cells due to their higher firing rates. Surprisingly, the model predicts that pyramidal cells (‘cADpyr’-firing type) have twice the positive influence on sorting accuracy compared to interneurons. This likely reflects their larger morphological footprint, which increases their visibility across multiple electrodes. Additionally, the model predicts higher sorting accuracy for neurons in layer 5 across both regimes and in layer 6 during spontaneous activity (Fig. 6e). Another bias emerges in the evoked regime, where delayed non-accommodating (‘dNAC’) neurons are more accurately sorted than other electrical types (Fig. 6f).

Taken together, these findings indicate that Kilosort4’s output is highly sensitive to neurons’ anatomical and physiological properties, which create systematic biases in sorting accuracy. The strongest biases should favor high-firing pyramidal neurons with large spatial spike footprints located in layer 5.

We investigated whether spike sorters exhibited selection biases, by over-representing certain model neuron types while undersampling others (Fig. 6g,h). To quantify these biases, we assigned each sorted ‘single-unit’ the biophysical features of its best-matched ground truth neuron and assessed whether neuron types were more or less likely to appear after spike sorting compared to their representation in the ground truth. Excitatory pyramidal neurons, which made up 91% of the ground truth population (Fig. 6g), were consistently under-represented across all layers by approximately a factor of 2, except in layer 5, where they were over-represented by the same factor (Fig. 6h, ratios in blue; Fisher’s exact test, two-tailed, *p* < 0.05). In contrast, most interneurons were proportionally represented relative to the ground truth, except for rare interneuron types, which were over-represented by factors ranging from 2.9 to 8.1 by some spike sorters, particularly Kilosort (1 to 2.5).

This selection bias, driven by kilosort4’s sensitivity to neurons’ feature, has substantial consequences: because high-firing neurons are preferentially retained, mean firing rates in most cortical layers are overestimated by a factor of three to four in sorted spike trains compared to ground truth (compare orange and blue distributions in Supplementary Fig. 7a, normalized; Supplementary Fig. 7b, raw, and the increasing ratio of sorted vs. true units for increasingly active units Supplementary Fig. 7c). The most pronounced overestimation occurs in layer 5 (Supplementary Fig. 7e), when excluding layer 1 (Supplementary Fig. 7d), where the low sorting yield prevents reliable conclusions.

These results demonstrate that the low yield of well-sorted units is driven by sensitivity to neuronal features, rather than an inherent limitation of sorting algorithms. However, this sensitivity also introduces distortions in estimated circuit activity, particularly systematic over-estimation of firing rates [12] (Supplementary Fig. 7).

#### Spike sorting degrades stimulus representation

After characterizing the different types of spike sorting biases, we found that these biases could significantly distort the cortical firing rate distribution. This distortion has the potential to lead to misinterpretations of neural coding principles and impair stimulus decoding from recorded cortical activity, since neural representations derived from extracellular recordings are only indirectly accessible to experimentalists through spike sorting.

To assess the extent of these distortions, we designed an experiment to quantify how spike sorting biases degrade stimulus representation. Specifically, we evaluated how much information was lost or altered in the sorted spike trains compared to the ground truth responses. We simulated cortical responses to whisker deflections repeated 50 times in eight different directions, spaced 45 degrees apart (Fig. 1d,g). These deflections were modeled by activating for 200 ms distinct configurations of thalamic fibers arranged in contiguous circular sector geometries (Fig. 1g). The population response to each stimulus was defined as the spike count of each neuron during stimulus presentation. To visualize the high-dimensional structure of these responses, we applied principal component analysis (PCA), chosen for its interpretability, ease of use, and efficiency, to reduce the data to three dimensions. This transformation provided a direct comparison between the geometry of the true neural representations (neural latents) and their sorted counterparts, revealing how sorting affects stimulus encoding. We then plotted the resulting neural manifolds (Fig. 7a), where each dot represents a trial, and colors indicate the different stimulus directions.

**Fig. 7.**
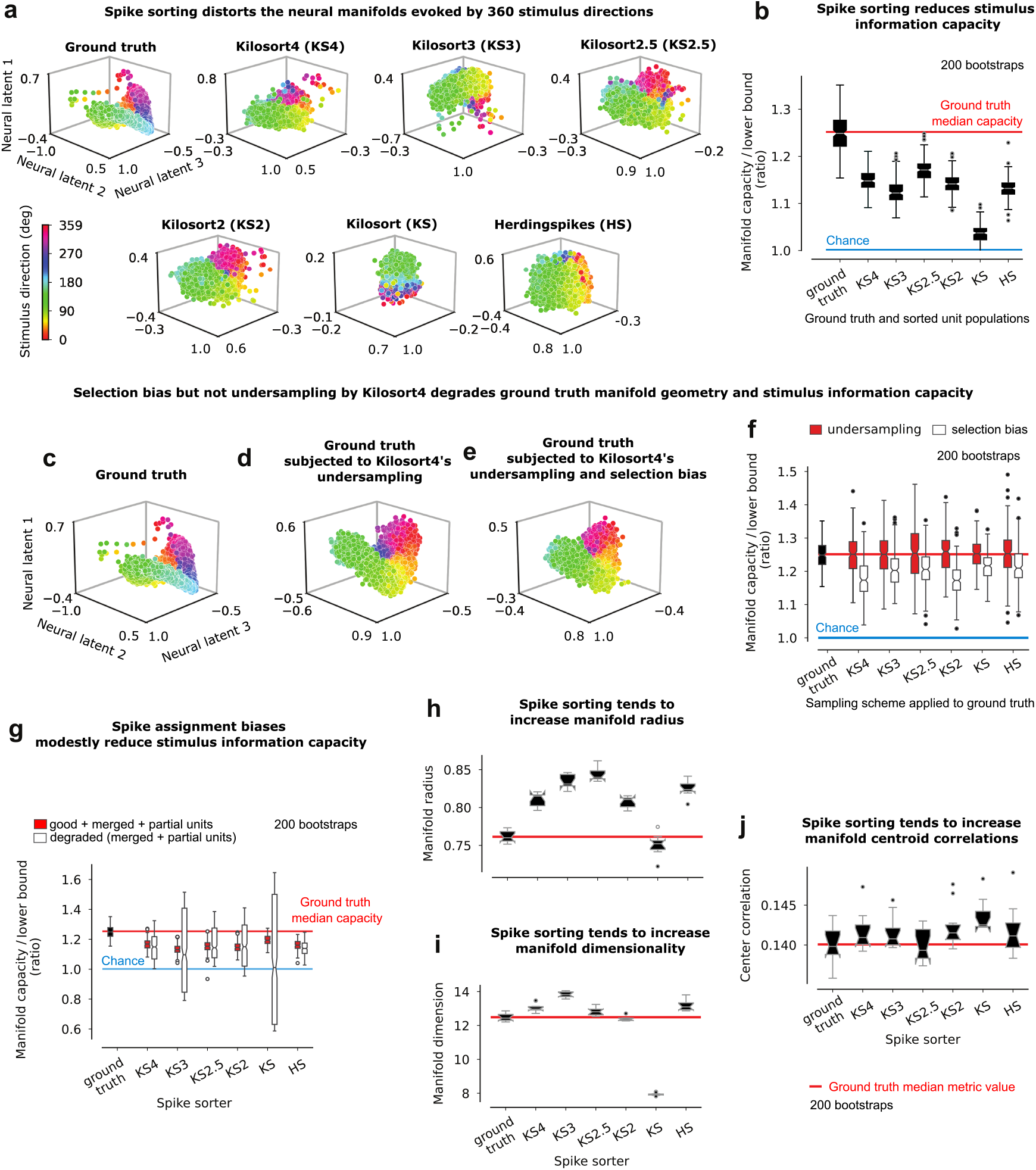
Impact of spike sorting biases on stimulus discriminability. **a**. Neural manifolds of ground truth and sorted populations. PCA was applied to neuron responses to 360 stimulus directions separated by 1 degree, a dense sampling of the stimulus space, to assess the continuity of stimulus mapping on the manifold. and **b**) information capacity. Information capacity distributions are obtained from 200 bootstraps of ground truth neurons and spike sorted units. Chance level is 1 when the manifold capacity equals the lower bound capacity calculated by shuffling the mapping between neural responses and stimulus directions. Neural manifolds of **c**) ground truth as in **a**, of **d**) ground truth subjected to Kilosort4’s undersampling bias by randomly undersampling ground truth neurons to match Kilosort4’s yield, **e**. ground truth subjected to Kilosort4’s undersampling and selection biases by sampling ground truth neurons to match both Kilosort4’s yield and its distribution of sorted unit types. **f**. Manifold information capacity for the undersampling **d** (red box) and selection biases in **e** (white box). Information capacity distributions are obtained from 200 bootstraps of ground truth neurons **g**. Information capacity for ground truth neurons, sorted units that both ‘good’, merged and partial (pink Fig. 5d) and degraded units (‘merged’ and ‘partial’) in Fig. 5c,d. The distribution of unit types were matched between the two. **h**. Manifold radius: average distance between the centroid and boundaries of the manifold evoked by the repetition of a stimulus direction (object manifold, [54, 55]), **i**. Manifold dimension: it is the average dimensionality of the embedding subspace spanned by these object manifolds ([54, 55]). **j**. Center correlation: correlation between object manifold centroids. Distributions were obtained with 200 bootstraps of the population responses. Analyses were performed on the 1-hour biophysical simulation of Neuropixels evoked regime sampled at 20 kHz.

The ground truth neural manifold exhibited a ring-like geometry (Fig. 7a), which continuously reflected changes in the stimulus direction across trials, similar to ring topologies evoked by circular stimuli in mice head-direction circuit [52] and visual cortical areas [53]. Spike sorting distorted the manifold’s geometry for all spike sorters, but the most recent sorters still preserved stimulus mapping (from Kilosort2 to 4).

To obtain a precise, quantitative, and interpretable measure of the information loss induced by spike sorting distortions, we used information capacity, a metric that quantifies how much stimulus-related information is stored per neural latent. By assessing how well a linear classifier distinguishes between stimulus-evoked responses based on their manifold geometry, information capacity provides a direct way to evaluate sorting-induced distortions [54].

Because information capacity depends on neuron population size, we reduced the dimensionality of both ground truth and sorted populations to 200 neural latents, following [54], thus ensuring comparability between conditions. To enable statistical comparisons, we bootstrapped population responses 200 times, using Gaussian Random Projection for dimensionality reduction. Compared to PCA, this method is exponentially faster and more robust to outliers, and thus better suited for large-scale resampling.

We then measured information capacity using Mean-Field Theoretic Manifold Analysis (MFTMA) [54–56] per boostrapped population and normalized it to a lower bound, determined by shuffling the mapping between neural responses and stimuli. This allowed direct comparisons of the average information capacity between sorting conditions and with chance-level performance. As expected, averaged normalized information capacity remained stable when increasing population dimensionality from 50 to 1,600 (Supplementary Fig. 8), confirming that 200 dimensions were sufficient.

Distortions in manifold geometry were associated with significant reductions in information capacity, averaging a 50% decrease across spike sorters, from 25% above the lower bound (Fig. 7b, t(199) ranging from -68 to -24, all *p* < 0.001, Student’s t-test). These results highlight the extent to which spike sorting errors alter the structure of neural representations and degrade stimulus information.

The observed loss in discrimination capacity was partially explained by selection biases for ground truth neuron types, but not by random undersampling of neurons. Undersampling ground truth neurons to match spike sorting yields neither distorted the manifold’s structure nor degraded its information capacity (example for Kilosort4, Fig. 7d vs. c, and all sorters in Fig. 7f, red boxes vs. red line, t(199) ranging from -0.71 to - 0.02, all *p* > 0.47, Student’s t-test). However, resampling the distribution of ground truth neuron types to match that of spike sorted units visibly degraded the manifold structure (Fig. 7e vs. c) and its information capacity by an averaged 21% across sorters (Fig. 7f, white boxes vs red line, t(199) ranging from -17 to -8, all *p* < 0.001, Student’s t-test).

We assessed the impact of spike assignment biases on the information capacity of spike-sorted populations by comparing degraded sorted units (‘merged’ and ‘partial’) with ‘good’ sorted units. Here, ‘good’ units included all units with sorting accuracy above 80%, encompassing units that were both ‘good’, ‘merged’, and ‘partial’ (Fig. 5d). The distribution of sorted unit types differed significantly between degraded and ‘good’ units across spike sorters (p < 0.001, Fisher exact test with Monte Carlo simulations). So to isolate the effect of spike assignment biases from selection biases on manifold information capacity, we resampled degraded units to match their unit-type distribution to that of ‘good’ units.

On average, the manifold capacity of ‘good’ units was 37% lower than ground truth across spike sorters (Fig. 7g, red boxes vs. red line, t(199) ranging from -0.31 to -15, all *p* < 0.001, Student’s t-test). Degraded units exhibited an even greater reduction in capacity, averaging a 12% decrease relative to ‘good’ units across most sorters (Kilosort4, 2.5, 1, and Herdingspikes). These results highlight the strong detrimental effects of selection biases and spike assignment biases on information capacity (Fig. 7g, red vs. white).

According to the framework proposed by [55], information capacity is governed by three key geometrical properties: manifold radius, manifold dimension, and inter-manifold correlation, with reductions in these measures generally leading to increased capacity. To assess the impact of spike sorting distortions on information capacity, we systematically quantified their effects on these three metrics. We defined the manifold radius as the average distance between the center and boundaries of the manifold evoked by 50 repetitions of a stimulus direction (analogous to the object manifold in [55]). The manifold dimension represents the average dimensionality of the embedding subspace spanned by these object manifolds, while inter-manifold correlation quantifies the similarity between their centroids.

The observed reductions in information capacity were primarily driven by an increase in manifold radius, higher dimensionality, and more correlated centroids. Across sorting algorithms, direction manifolds’ radii increased by an average of 6%, except for Kilosort, which showed a 3% decrease (Fig. 7h, t(199) ranging from -87 to 0.18, all *p* < 0.001, Student’s t-test). Similarly, manifold dimensionality increased by an average of 6% across sorters, except for Kilosort and Kilosort2, which exhibited a 19% decrease (Fig. 7i, t(199) ranging from -59 to 234, all *p* < 0.001). Additionally, Kilosort3, Kilosort2, Kilosort1, and Herdingspikes showed a small but significant 1% increase in centroid correlation (Fig. 7j). These changes suggest greater overlap in neural responses to distinct stimuli, reducing the linear separability of stimulus-specific representations.

## 3 Methods

### 3.1 Datasets and simulations

#### Marques-Smith dataset

We used a 26 minute recording performed with the Neuropixels 1.0 Phase3A probe [4] in the primary somatosensory cortex of anesthetized adult rats aged between six weeks and eight months [8]. The probe, consisting of four columns with 96 electrode sites each, and rows spaced 20 µm apart, was aligned with a cortical column and spanned all six layers. The first electrode was positioned at the Pial surface. We used the cortical layer border depths from the Pial surface identified in P23-36 adult rats in [8, 57] to determine that layer 5 contained the highest number of isolated units. Traces were acquired at a sampling rate of 30 kHz, and analyses were performed on file c26, accessible at google-drive. The publicly shared recording underwent minimal processing, limited to offset subtraction [8].

#### Horvath dataset

We utilized a dataset from a single rat, which included recordings at three cortical depths with a single probe insertion [41]. Electric field potentials were recorded at a sampling rate of 20 kHz using an 8 mm long and 100 µm wide silicon probe with 128 dense recording sites, arranged in a 32 × 4 grid of square electrodes. Like Neuropixels 1.0, the electrodes were square-shaped and made from titanium nitride, with low impedance (50 kOhms) in saline, measured with 1 nA sinusoidal currents at 1 kHz, compared to Neuropixels 1.0’s 149 pm6kOhms [4]. Unlike Neuropixels 1.0’s staggered electrode layout, Horvath’s probe features equidistant electrodes. The electrodes are also larger (20 µm per side, compared to Neuropixels 1.0’s 12 µm), and electrode and positioned closer together, with a center-to-center distance of 22.5 µm for Horvath versus 25 µm for Neuropixels 1.0. We accurately reproduced the site locations from the experiment in [41] (Supplementary Fig. 1b). The dense probe captured activity from layers 1 and 2/3 at insertion depth 1, layers 2 to 5 at depth 2, and layer 6 at depth 3.

#### Synthetic model

We analyzed simulated Neuropixels recordings from [16], sampled at 32 kHz. The simulation included spiking activity from 250 units in layer 5, comprising 200 excitatory and 50 inhibitory biophysically simulated units with spike activity based on data from [30]. Importantly, this model lacks the cortical microcircuit connectivity that drives spiking dynamics and correlation statistics, whereas our proposed biophysical model includes these connections.

#### Biophysical model

We used a model of the rat primary somatosensory cortex, at age P14 [28]. Of the 4,234,929 neurons in the circuit, we simulated the voltage activity of a microcircuit column comprising 30,190 neurons using the NEURON software [58], via Blue Brain Project’s Neurodamus interface, [59] under spontaneous and stimulus-evoked conditions. We detected spikes from 1,388 model neurons during the spontaneous activity condition, 1,836 model neurons during the 1-hour evoked condition sampled at 20 kHz, and 1,552 neurons during the 10 minute evoked condition sampled at 40 kHz, within 50 µm of the Neuropixels probe. The network was calibrated to emulate an *in vivo* state of spontaneous firing with low synaptic release probability and asynchronous firing at low rates [29]. All model neurons fired at least once in the first 10 minutes, making detection theoretically possible. Meta-parameters included extracellular calcium concentration (modulating synaptic reliability), noise in external input compensation, and the overall activity level, described as a proportion of layer-specific reference firing rates. The model was configured with parameters P_fr_ = 0.3, [Ca] = 1.05, and ROU = 0.4 [29].

#### Probe reconstruction

We simulated extracellular recordings from 384 recording sites of a Neuropixels 1.0 probe, without accounting for electrode impedance, capacitance, or the effects of electrode shape and size. Instead, the electrodes were modeled as infinitesimal points [39]. For the Horvath setup, extracellular recordings were simulated from 128 electrodes, and the probe was positioned at three different depths in the cortical microcircuit to replicate the three in vivo locations [41]. However, the geometry of our microcircuit prevented us from perfectly matching the electrode distribution across layers observed in vivo (Supplementary Fig. 1b, Depth 1, Depth 3 had 68 sites in L6, 60 outside the cortex).

#### Generative model of the electric field

Neurons are modeled as multicompartment cables using the NEURON simulation environment [60]. Extracellular signals are calculated at simulation-time using the BlueRecording pipeline [39]. Briefly, we assume that the extracellular medium can be treated as homogeneous, with a conductivity σ, and effectively infinite relative to the size of the neural circuit and recording electrodes. We also assume that the recording electrode contacts can be treated as infinitesimally small.

At each time step, the simulator calculates the extra-cellular signal 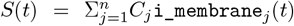, where i_membrane is the total compartmental transmembrane current, j is the index of each neural compartment in the circuit, and C is a precomputed scaling factor.

For simulations of the probes used by [41], we calculate the scaling factors C_j_ using the line-source approximation [40]. We assume that each neural compartment can be treated as a 1-dimensional line.

For each neural segment, the scaling factor 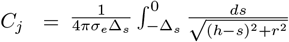, where Δ_s_ is the length of the segment, h is the distance from the near end of the segment to the recording electrode in the axial direction, and r is the absolute value of the distance from the near end of the electrode to the electrode in the perpendicular direction.

For simulations of Neuropixels probes, we calculate the scaling factors C_j_ using the point-source approximation. We assume that each neural compartment is infinitesimally small. The scaling factors 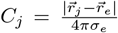, where 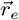 is the position of the electrode and 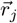 is the position of the compartment.

#### Sampling frequency

The simulated Neuropixels traces were sampled at 40 kHz in the spontaneous regime and at 20 and 40 kHz in the 1-hour long and 10 minute evoked regimes respectively which are close to sampling frequencies used in past studies [9, 16, 41, 61]. The traces collected with the reconstructed Horvath probe were sampled at a frequency of 20 kHz as in [41].

#### Fitting amplitude and background noise

The biophysical traces missed noise from external sources and raw voltage amplitudes ranged from − 1.26 µV to 0.6 µV, a narrow range near zero that reduced spike resolution, causing their omission after Kilosort processing, as it assumes 16-bit integers. To address these, we scaled the traces and modeled external background noise as an additive Gaussian distribution with zero mean across all electrodes, following [7, 16]. We ensured biological validity by fitting the trace extrema and background noise to Marques-Smith data in three steps: applying a gain derived from dividing the extrema of preprocessed *in vivo* and simulated traces, adjusting noise levels measured by the trace mean absolute deviation (MAD), for each layer, and manually fine-tuning the gain and noise after preprocessing. All fits were performed using the Nelder-Mead algorithm. We used the same gain and noise parameters for both spontaneous and evoked regimes. The best-fit gain was 677, with missing noise root mean square values of 2.8 µV, 3.4 µV, 3.7 µV, 3.6 µV, and 3.8 µV for layers 1, 2/3, 4, 5, and 6. The spontaneous regime’s maximal amplitude (361 µV) closely matched Marques-Smith’s (362 µV), while the evoked regime produced a slightly higher value (387 µV). The gain to fit the synthetic traces’ extrema to Marques-Smith’s data was 0.41.

For the denser probe, separate gains were derived for each depth, and background noise was fitted to Horvath data [41] by matching depth and layer. Gains were 12,463, 1,949, and 20,778 for depths 1, 2, and 3, respectively. Missing noise RMS values were 27.3 µV and 30.3 µV for layers 1 and 2/3 at depth 1, 37.4 µV and 37.9 µV for layers 4 and 5 at depth 2, and 35.6 µV for layer 6 at depth 3. The maximum amplitudes closely matched Horvath data, with errors below 3.5%.

#### Preprocessing

For all analyses, the traces were high-pass filtered above 300 Hz as in [4, 6, 7, 9, 16], using *SpikeInterface* [16] to attenuate their low-frequency noise component while preserving spikes’ frequency band. The common median voltage across electrodes was then subtracted to remove the global noise component shared across electrodes (common median referencing [62]). Whitening was applied by all Kilosort sorters to reduce local spatial correlations across channels [7].

### 3.2 Model validation

#### Amplitude-to-noise ratio

To minimize computational cost, we analyzed the first 10 minutes of all recordings from cortical electrode sites. The amplitude-to-noise ratio (ANR) was calculated by dividing the trace amplitudes by the median absolute deviation (MAD). We compared ANR distributions across experiments by using the largest ANR range, divided into 100 bins, and calculated the probability of each ANR bin per electrode and experiment. The ANR was then averaged across electrodes and plotted with confidence intervals across electrodes for each experiment (Fig. 2q).

#### Spectral composition

Power spectral densities (PSD) were computed with the Welch method [63], at 1 Hz resolution with the Scipy package. Filtering window sizes matched recording sampling frequencies, and PSDs were denoised with a Hann window with a one-second overlap.

Power law fits were performed by fitting a linear regression to the log-transformed PSDs, whose coefficients were the α parameters of the 1/f^α^ power laws.

#### WaveMap

Single-unit spikes were extracted using spike-triggered averaging. The six nearest electrode sites, which exhibited the largest average spike amplitudes, were selected and ranked in descending order based on the first positive peak, trough (maximum negative amplitude), or second positive peak—depending on which had the largest absolute amplitude. Spikes were aligned at the trough over a 12 ms window centered on the spike timestamp, later reduced to 6 ms. Waveform amplitudes were normalized to a range of 0 to 1, and UMAP (umap-learn v0.5.3) was applied to project the waveforms into a two-dimensional latent space. The resulting embedding was clustered using the Louvain algorithm (python-louvain v0.16) as in [9, 47].

Spike-triggered average of isolated intracellular recordings were calculated for the active morpho-electric types that were found within 50 µm of the Neuropixels electrodes (Supplementary Fig. 3).

#### Firing rate

Since firing rate distributions followed a lognormal pattern, we compared simulated and *in vivo* firing rates using their decimal logarithm (log) transformations. A Mann-Whitney U test was conducted to assess whether the medians and variances of the log-transformed firing rates of the sorted units were similar (Fig. 4b). To calculate the median and confidence intervals of the variance shown in Fig. 4b, we bootstrapped the firing rates 100 times to generate a distribution of variances.

### 3.3 Spike sorting

Spike sorting was performed on the probe-wired raw recordings using SpikeInterface version 0.100.5 [16]. Sorting generally succeeded with default parameters. For all sorters, we used the default settings, with a few key adjustments: the minimum firing rates for unit detection (*minFR*) and site curation (*minfr_goodchannels*) were both set to 0 to maximize unit detection. This adjustment enabled a fair comparison with sorters like Kilosort 4 and Herdingspikes, which do not filter units based on firing rates. Additionally, batch sizes were adjusted to accommodate GPU memory constraints.

We report results from Kilosort2 for 4 minutes of evoked recordings, as it failed to process longer durations. Kilosort2.5 only functioned without drift correction (with the *nblocks* parameter set to 0). Mountainsort 4 was not used due to its long sorting time, which exceeded eight hours. Klusta and HDSort were excluded because they are not maintained to be compatible with Python versions later than 3.9.

All tested sorters, except for the original Kilosort version and Herdingspikes, isolated ‘single-units’ and labeled them as either ‘good’ or ‘multi-unit’ in the *KSLabel* metadata. For Kilosort and Herdingspikes, all sorted units were reported.

### 3.4 Sorted unit quality

#### Unit isolation ratio

Kilosort (2, 2.5, 3, 4) isolate ‘single-units’ as clusters of spikes with minimal violations of the neuronal refractory period — a brief interval following a spike (1–5 ms [7]) during which neurons cannot generate another spike. Units that do not satisfy this criterion are classified as ‘multi-units’. To distinguish between these categories, these sorters calculate the “contamination rate”, which measures the proportion of refractory period violations.

This metric is defined as the ratio of the frequency of spikes occurring within the refractory period to the baseline frequency of spikes outside this period [64]. A contamination rate below 0.1 is used as the threshold for identifying ‘single-units’ [7]. The unit isolation ratio was calculated as the proportion of ‘single-unit’ to the ‘multi-unit’ produced by the spike sorters. To investigate the impact of waveform duration on unit isolation, we ran Kilosort4 on a biophysical simulation of the spontaneous regime, processing the data at twice the normal speed. This was achieved by downsampling the trace to every second sample while maintaining the original sampling frequency of 40 kHz, as described in [7].

#### Unit yield

The sorting yield refers to the number of ‘single-units’ isolated by Kilosort versions 2.0, 2.5, 3.0, and 4.0. This measure of sorting quality is accessible to experimentalists who do not have access to ground truth data. To enable comparisons with *in vivo* studies and across experiments with varying cortical column coverage, we employed the yield per site metric.

#### Agreement Score

We utilized the agreement score defined in [16], which is the ratio of correct spike coincidences between a ground truth neurons and a sorted unit to the total number of spikes.

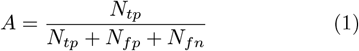

where *N*_*tp*_ is the number of true positives (detected spikes), *N*_*fp*_ is the number of false positives (spurious spikes), and *N*_*fn*_ is the number of false negatives (missed spikes). A score of one hundred percent can only be achieved when all spikes from the ground truth and the sorted units coincide.

Coincidences were detected within a time window of Δ_*t*_ = 1.3 ms around each ground truth spike. This duration maximized sorting accuracy for Kilosort4 (Supplementary Fig. 6a). To ensure that a Δ_*t*_ of 1.3 ms was not overly lenient — potentially assigning high scores for random matches, we compared the distribution of ground truth neuron sorting accuracies for zero shift versus long constant shifts of ground truth spike timestamps. When Δ_*t*_ becomes too long, the accuracy distribution will show a single peak near 0, indicating chance-level performance. For meaningful Δ_*t*_, the distribution will display two peaks: one for correctly detected units and another near chance level for missed units.

In our simulation, a spike waveform is timestamped when the voltage reaches − 10 µV, while spike sorters may timestamp the spike at a different point in the spike waveform. As we expected, for small shifts, the accuracy distribution for ground truth neurons showed two distinct peaks. However, with larger shifts (e.g., 30 ms), the accuracies degraded and peaked at 0, indicating chance-level performance (Supplementary Fig. 6b). Based on this analysis, we finalized Δ_*t*_ = 1.3 ms with no shift.

Coincidences between two independent spike trains can occur by chance, resulting in non-zero agreement scores. We assumed that the sorted unit’s spikes occurred independently and stochastically with a constant firing rate ν. Under this assumption, the probability of observing at least one coincidence (a true positive tp) within the interval surrounding the ground truth spike (T = 2 ∗ Δ_*t*_) is given by the homogeneous Poisson equation:

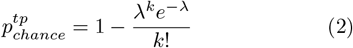

where k = 0 and the term 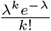 is the probability of observing no coincidence. λ is the expected number of coincidences within the interval T and for a firing rate *ν* = min(*N*_*true*_, *N*_*sorted*_) taken as the firing rate of the least active unit between the ground truth and the sorted unit:

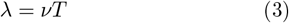

Chance agreement scores *A*_*chance*_ were calculated following equation 1 for each pair of ground truth neuron and sorted units by setting the number of true positives to 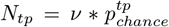, the number of false positives to *N*_*tp*_ = *N*_*sorted*_ − *N*_*tp*_ and the number of false negatives to *N*_*true*_ − *N*_*tp*_.

#### Sorting accuracy

We defined the sorting accuracy of a ground truth neuron as the highest agreement score achieved with its best-matching sorted unit.

#### Spike waveform quality metrics

To obtain a full quantitative description of sorted ‘single-unit’ spiking, we used three complementary categories of metrics: 1) spike waveform quality (SD ratio, MAD ratio, and SNR), 2) unit isolation quality (silhouette score, ISI violations ratio, refractory period contamination metrics, amplitude cutoff) and 3) firing statistics (firing rate and firing range). Except for the MAD ratio, all metrics were calculated using SpikeInterface [16] and have been introduced in previous studies [20, 65–67]. We did not include computationally expensive metrics such as isolation distance or nearest-neighbor measures, nor did we analyze drift metrics, since our simulated recordings assume stationary electrodes without drift.

To assess waveform consistency, we used the SD ratio [65], which measures the ratio of the standard deviation of a unit’s spike trough (negative peak) to the standard deviation of background noise. This metric approaches 1 for well-isolated single neurons but increases with contamination. We introduced a new, more outlier-robust metric, the MAD ratio, which replaces standard deviation with mean absolute deviation:

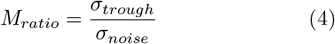

We calculated units signal-to-noise ratio (SNR) to assess their isolation from background noise, a key factor in identifying well-isolated units. This was done by identifying the electrode site where the unit’s average spike displayed the largest trough. The spike’s extremum (maximum/minimum voltage) was then measured and divided by the median absolute deviation of the background noise from that electrode. A low SNR value (close to 0) indicates high contamination by noise, suggesting poor unit isolation.

#### Unit isolation quality metrics

To evaluate unit isolation, we used the ISI violations ratio (isi_violations_ratio), a key metric for detecting contamination based on the fundamental principle that neurons exhibit a refractory period preventing them from firing consecutive spikes within a short interval. This ratio quantifies the proportion of a unit’s spikes that violate this period (set at 1.5 ms), indicating potential contamination from other units. To complement this, we also computed two related measures: 1) refractory period contamination rate (rp_contamination), the proportion of contaminant spikes violating a unit’s refractory period, and 2) refractory period violations count (rp_violations), the absolute number of these violations [67]. These three ISI-based metrics collectively provide a comprehensive, quantitative characterization of contamination, capturing both the degree of contamination of a unit relative to its level of activity and the magnitude of the contamination from other units.

The silhouette score, a widely used clustering metric, quantifies how well a unit’s spike waveforms are separated from other clusters. A silhouette score near 1 indicates compact, well-separated clusters, while a score near -1 suggests poor isolation [66].

The amplitude cutoff metric estimates the completeness of a unit by measuring the fraction of spikes with amplitudes near the upper limit of the unit’s spike amplitude histogram. Assuming that a unit’s spikes are symmetrically distributed, a high amplitude cutoff suggests missing spikes [16]. We used raw spike amplitudes instead of the template scaling factors generated by Kilosort sorters.

#### Firing quality metrics

To quantify firing rate variability, we measured the firing range, calculated as the difference between the 95th and 5th percentile firing rates over short time bins (e.g., 10 s). Since our simulated recordings assume stationary electrodes without drift, unphysiologically high firing ranges may indicate noise contamination rather than real neuronal activity. We also included firing rate in our analysis.

#### ‘Single-unit’ curation

The logistic regression model was a generalized linear model for the Binomial family and the logit link (statsmodel). The models’ precisions and recalls for more than 80% accurate sorted single-units (‘good’ units) were calculated by splitting the dataset into an 80% train set and a 20% test set randomly sampled a hundred times (number of folds). We selected the unit quality metrics that were the most efficient to compute and that could be computed for the largest number of units (n=182 units) to maximize the size of the training dataset. The quality metrics were z-scored before training and testing, which allowed to compare their weights from the fractional logistic model. Z-scoring did not noticeably change the precision and recalls of the models.

CEBRA is an autoencoder with robust generalization performance conferred by contrastive learning optimization of the InfoNCE loss. We used 40 iterations which were sufficient for the convergence of the loss functions for all cross-validation groups of samples (the folds). We used the smallest possible number of iterations to minimize the large computational cost of running 100-fold cross-validation. We manually searched for the best set of hyperparameters, by changing one parameter at a time. We changed the number of spike waveforms (25 and 50) in the dataset, CEBRA’s output dimensions (3 and 10) and the k-NN’s number of neighbors (1, 2 and 10, 20). While a more exhaustive parameter search and regularization can improve the model performance, we favor logistic regression for its efficiency and interpretability.

#### Model of sorting accuracy

We trained a generalized linear model for the Binomial family and a logit link (statsmodel v0.14.2) to predict model neuron sorting accuracy based on 21 available descriptive features. The model weights were regularized with Lasso (L1-norm) which enables feature selection by setting the weights of linearly dependent unit features with high variance, which effect on sorting accuracy can not be disambiguated, to zero. We ensured model convergence by increasing the number of iterations (maxiter parameter) and reducing the loss function convergence tolerance (cnvrg_tol) until the models’ cross-validated explained variances (*R*^2^) became stable. The models converged within 100 iterations with a convergence tolerance of 1e-10.

### 3.5 Neural manifold information metrics

We used information metrics to quantify the impact of the identified biases on the linear separability of sorted cortical circuit responses evoked by simulated whisker deflection stimuli. To manage the computational cost, the metrics were calculated for eight stimulus directions, separated by 45 degrees, out of the 360 tested (coarse discrimination task). The 360 whisker deflection stimulus directions were simulated using 200 ms thalamic fiber activations arranged in contiguous circular sectors, with each direction repeated continuously 50 times.

We conducted a mean-field theoretic analysis using the neural_manifolds_replicaMFT package, available on [Github](https://github.com/schung039/neural_manifolds_replicaMFT), to calculate the average manifold capacity, radius, and dimensionality for each category of spike sorting bias [54]. Dimensionality reduction was applied to all datasets to ensure the same number of unit features, set to 200 (the default parameter n_t). Capacity was then normalized by dividing it by the lower bound capacity of each dataset, which can be empirically calculated from a shuffled label dataset. Medians and confidence intervals were obtained by resampling each dataset 1,000 times. The model’s number of samples n_t was set to 200 (its default value), and the margin κ was set to zero.

### 3.6 Statistical tests

Kruskal-Wallis H-tests were performed to assess whether trace median peak amplitude, noise (M: n=10, 42, 34, 42, 98 sites for layers 1, 2/3, 4, 5 and 6 respectively; NS and E: 16, 47, 19, 52, 68 sites; H: 36, 60, 20, 88, 68 sites; DS: 37, 64, 32, 88, 76 sites), AP power and frequency scaling slope changed across layers (Fig. 2). It is a non-parametric test that tests the null hypothesis that the population median of all of the groups are equal. Measurements were taken from distinct samples. Mann-Whitney U tests were used per layer to compare median peak amplitudes between models and *in vivo* data. They were also used per layer to compare the median decimal logarithm of the firing rates and the variance of the decimal logarithm of the firing rates (Fig. 4). A Pearson’s χ^2^ goodness-of-fit test was used to determine whether proportions of ‘single-units’ were different from the expected (i.e., average) proportions across experiments (Fig. 5). It was also used to determine whether proportions of ‘good’ neurons, that is, of neurons sorted with above 80% sorting accuracy, were different from expected proportion across spike sorters (Fig. 6b). This test is appropriate as all sample sizes were larger than 5. All three tests were implemented with the Scipy package.

We measured the explained variance of the fractional logistic regression using McFadden’s pseudo *R*^2^ (Fig. 6c and d), as it provides a more appropriate definition of the log-likelihood for the model based on the observed data in logistic regression, than Pearson correlation. Student’s t-tests were used to test the null hypothesis that the logistic and fractional logistic regression model weights were statistically different from zero. Statsmodel was used to calculate the model’s log-likehood, the null hypothesis’ necessary to calculate the model pseudo *R*^2^, and the Student’s t-tests.

We used a Fisher exact test of goodness-of-fit, rather than a χ^2^ test, to determine whether the probability of occurrence of each model neuron type was the same in the ground truth and the population of spike-sorted units (Fig. 6h), as some samples were smaller than 5.

Due to the computational intensity of the Fisher exact test, we used custom codes of Fisher Monte Carlo test, which we first validated against the Fisher exact test for small contingency tables. It was also used to test whether unit-type distributions among degraded units and ‘good’ units was the same.

### 3.7 Software and hardware

Extracellular signals were calculated using an experimental version of the BlueRecording [39] pipeline. Source code and instructions for precomputing the scaling factors for each compartment are available on GitHub. Analyses were programmed in Python v3.9.7 and Matlab R2022b. The Fractional regression model was implemented with Statsmodel in Python. All Kilosort sorters were run with Matlab with the Parallel Computing Toolbox, Signal Processing Toolbox and the Statistics and Machine Learning Toolbox. Software versions for all analyses were Spikeinterface: 0.100.5 ([16]), Statsmodel: v0.14.3,, Scipy: v1.13.1, Matlab’s Parallel Computing Toolbox: v7.7, Signal Processing Toolbox; v9.1, Statistics and Machine Learning Toolbox: v12.4, The codebase for the analyses is available on GitHub.

We run simulations parallelized on 120 cpu clx nodes (384 GB of memory: 12 DIMMs (6 per CPU) of each 32GB at 2933MT/s 2 Intel Xeon Gold 6248 CPUs at 2.50GHz with max speed of 3.9Ghz). Each CPU has 20 physical cores. Due to hyperthreading, slurm provides 80 cores per node scheduled with Slurm. With this setup a simulation of 10 minute of extracellular recordings typically took 6 days.

Spike sorting was supported by GPUs (4 volta v100 skl each with 16GB of RAM) on nodes with 768 GB of memory: 12 DIMMs (6 per CPU) of each 64GB at 2666MT/s, 2 Intel Xeon Gold 6140 CPUs at 2.30GHz with max speed of 3.7Ghz). Each CPU has 18 physical cores => 36 physical cores in total. Due to hyperthreading, slurm provides 72 cores per node.

Biophysical simulations were performed on a computer cluster with resource allocation managed by SLURM and Python multiprocessing framework was used to speed up preprocessing and data analysis.

### 3.8 Data availability

The dataset will be publicly available upon publication.

## 4 Discussion

We simulated dense recordings from a large-scale, detailed model of the rat cortical microcircuit designed to capture neuron heterogeneity in a data-driven and biophysically-detailed manner. Our goal was to test if spike sorting sensitivity to neuronal biological features could explain the large gap between the yield of modern spike sorters and the theoretical yield in a data-driven and biophysically-principled manner and characterize the theoretical impact of this. We found that a fractional regression model, where several model neuron features strongly impacted spike sorting accuracy, effectively accounted for the proportion of spikes correctly detected and assigned to isolated units. The model accounts for the yield gap by predicting reduced detection and isolation of neurons within isolable distances, influenced by the combination of electrode distance, layer-specific location, morphology, synaptic and electrical properties. The model further predicts that most presumably isolated units are sorted with accuracies below 80%, include spikes from multiple neurons and/or are incomplete and that all spike sorters display selection biases for particular unit types. Yet despite these biases, the most recent spike sorters preserved the overall low-dimensional shape of stimulus-evoked responses as well as most stimulus information content.

In agreement with evaluations reporting sorting accuracies above 80% on synthetic datasets [7, 16, 21], most tested spike sorters achieved high yields, accurately assigning spikes to over 85% of the neurons in synthetic datasets. However, in contrast to this predictions but in agreement with reports from dense extracellular recording experiments [4, 8–11], the spike sorters isolated about 200 neurons on the biophysical models — only a tenth of the achievable yield predicted by theoretical models [1, 12]. This comparison between synthetic model yield and theoretical yield was enabled by the scale and high level of biophysical detail of the model. It closely reproduces the average neuron density distribution across the full thickness of the rat primary somatosensory cortex [30]. The cortical microcircuit comprises 30,000 morphologically detailed neurons, captures cortical neuron diversity in 60 morphological types, grouped into inhibitory and excitatory synaptic types and 11 electrical neuron types, spanning all six cortical layers Figand receiving inputs from simulated thalamic fibers, which have been extensively validated against *in vivo* data [28– 30]. It also features realistic synaptic connectivity and short-term plasticity which constrain firing synchrony, critical for spike sorting [18], and are not modeled by current synthetic datasets. The model is also grounded in a biophysically principled description of spike amplitude decay with distance [38, 39].

We demonstrate that the extracellular traces simulated from our detailed model replicate key characteristics of two published experimental datasets from the rat primary somatosensory cortex [8, 41]: they exhibited spiking components, spectral composition, amplitude-to-noise ratios that were similar and/or in the *in vivo* range of these datasets. They contained spikes with five temporal profiles that matched those reported in these [8] and other studies in humans [9] and cats [46] and the dependencies on electrode distance and location reported in yet another set of studies [1, 12, 45]. Extracellular spike amplitudes remained resolvable within 75 microns and disappeared by 200 microns, in line with prior research [1] and ruling out the possibility that the low yield is attributed to a non-realistic rapid decay of spike amplitude with distance. The spike power’s tendency to increase with cortical depth may come to further validate the model’s accurate description of the distribution of neuron density and activity level across layers, as spike power has been shown to reflect neuronal density and activity across cortical thickness [42].

The biophysical models did not replicate the slow oscillations observed under anesthesia with the current parametrization; however, they are capable of generating activity resembling other brain states under alternative meta-parameter combinations. This flexibility makes them suitable for quantifying the impact of different brain states on the quality of spike sorting in the cortex [68].

Although the biophysical simulations produced spike waveforms that were longer than those observed *in vivo* — a factor known to potentially degrade spike sorting accuracy and increase false positives [7] — the resulting accuracy losses were minimal. We further demonstrated that reducing the waveform length by half through sub-sampling the traces, as suggested by Pachitariu et al. [7], did not significantly alter the number or proportion of isolated ‘single-units’. Additionally, for the non-subsampled traces the unit yield was already statistically consistent with the Marques-Smith dataset [8]. Finally, Pachitariu et al. [7] reported false positives as the main error resulting from longer waveforms, while we observed very few false positives in our results.

We developed a fractional regression model that succinctly describes and quantifies the relationship between cortical neuron location, biological features, and the proportion of correctly isolated spikes. Our findings reveal that, aside from distance, a sparse firing rate is the strongest determinant of low sorting accuracy, corroborating other studies that highlight the dependence of sorting accuracy on firing rate [7, 16]. This also explains previous experimental evidence showing that spike activity levels estimated from extracellular recordings can sometimes be significantly higher than the true average level of spiking activity measured with intracellular recordings [12] and predicted by modeling studies [69], as extracellular methods shift firing rate distributions toward higher values (Supplementary Fig. 7). This has an immediate impact on modeling work that often uses sorted firing rates as targets for their simulations [24, 25, 70]. It could be argued that the presence of low-firing rate neurons in our simulations, and hence this conclusion, is artificial. We note that, if the low-firing units were spiking at higher frequencies they would be more likely to be detected by the sorters, increasing the yield. The yield observed for our simulations is already higher than for the *in vivo* traces.

Additionally, our model identifies the second largest factor impacting sorting accuracy as the spatial footprint of spikes, aligning with analyses on spike spatial extent [7]. This finding directly supports recent demonstrations that denser probes than Neuropixels could enhance spike sorting yield threefold in cortical recordings by capturing small-footprint neurons more effectively [6] and is consistent with the observation that approximately half of isolated units with Neuropixels probes span multiple electrodes [9]. The fractional regression model offers a concise framework that disentangles and quantifies the effects of the key factors hypothesized to influence spike sorting accuracy [1, 12].

A novel and important prediction of the model is that a significant portion of the units isolated by the tested sorters are merged and incomplete, with sorting accuracies below 80%. The primary quality metric used to distinguish ‘single-units’ from ‘multi-units’ is the inter-spike interval (ISI, [7]). This metric is based on the reasoning that all neurons exhibit an absolute refractory period; thus, any cluster with a high proportion of ISIs less than 1–2 ms cannot represent a well-isolated unit. However, the opposite does not hold true: a clear refractory period does not necessarily indicate good isolation quality. For instance, a cluster containing intermixed spikes from two different neurons firing at distinct times may still show a clear refractory period.

To address this issue, we propose a new, straightforward model based on a linear combination of six quality metrics [16]. We demonstrate that this model effectively differentiate high-quality ‘single-units’ from merged and incomplete units among those produced by the Kilosort sorters. The model’s principles are intuitive: merged and incomplete units typically exhibit lower firing rates, a wider range of firing rates [16], greater shape variation (quantified by the median absolute deviation ratio, or ‘mad_ratio’ which we introduce here), and clusters that are more cohesive and better separated from others, as indicated by a silhouette score close to 1 [71].

In addition to these spike assignment biases, spike sorters displayed similar selection biases, particularly for excitatory pyramidal cells. Pyramidal cells were under-represented in most layers, and overrepresented in layer 5 across sorters. Increased activity evoked by the model stimuli in Layer 5, which is an important input layer, receiving inputs from our model thalamus may contribute to the over-representation of excitatory cells in layer 5 compared to other layers. Why layer 2/3, which is also an input layer, did not show such an increase in excitatory cells could be explained by different morphologies of the cells in these two layers. Further analyses will also be needed to determine why a few rare interneurons were over-represented by some spike sorters while most were accurately represented. Altogether, these findings suggest that most selection biases may stem from mechanisms shared among sorters, while others are influenced by processes unique to each spike sorter. Understanding these characterized selection biases is crucial, as recording from statistically representative samples of isolated neurons is a high-priority objective in systems neuroscience [1]. As combined Neuropixels and optogenetic techniques soon become available, it will be possible to validate these predictions.

## 5 Conclusion

This study underscores the critical limitations of current spike sorting methods in capturing the full complexity of dense cortical activity, revealing significant undersampling, assignment, and selection biases that impair accurate neural representation. By leveraging a biophysically-detailed model of the rat cortical column, we have demonstrated that current algorithms capture only a fraction of isolable neurons and preferentially sort certain neuron types, thus distorting stimulus encoding. Despite undersampling not directly diminishing information capacity, biases in neuron selection and assignment notably reduced stimulus discriminability, affecting the interpretation of cortical responses. These findings highlight the value of realistic, high-resolution models as complementary tools for improving spike sorting algorithms, setting a path toward more accurate large-scale neural activity representations that can deepen our understanding of cortical function and the neural basis of behavior.

## Acknowledgements

The authors would like to thank Enny Van Beest, Jennifer Colonell, Alessio Buccino and Marius Pachitariu for very insightful and critical feedback; Enny Van Beest, Ilias Rentzeperis and Cameron Mckenzie for helpful discussions and paper review suggestion; Alessio Buccino, Daniela Egas Santander, Christoph Pokorny, Andras Ecker, Natali Barros Zulaica for helpful discussions and support in identifying and accessing data analytical resources; the BBP Core Services team for responding to IT requests and services surrounding the research; Karin Holm for support of manuscript development and helpful discussions.

## Declarations

The authors declare no competing interests.

## Supplementary figures

**Supplementary Fig. 1.**
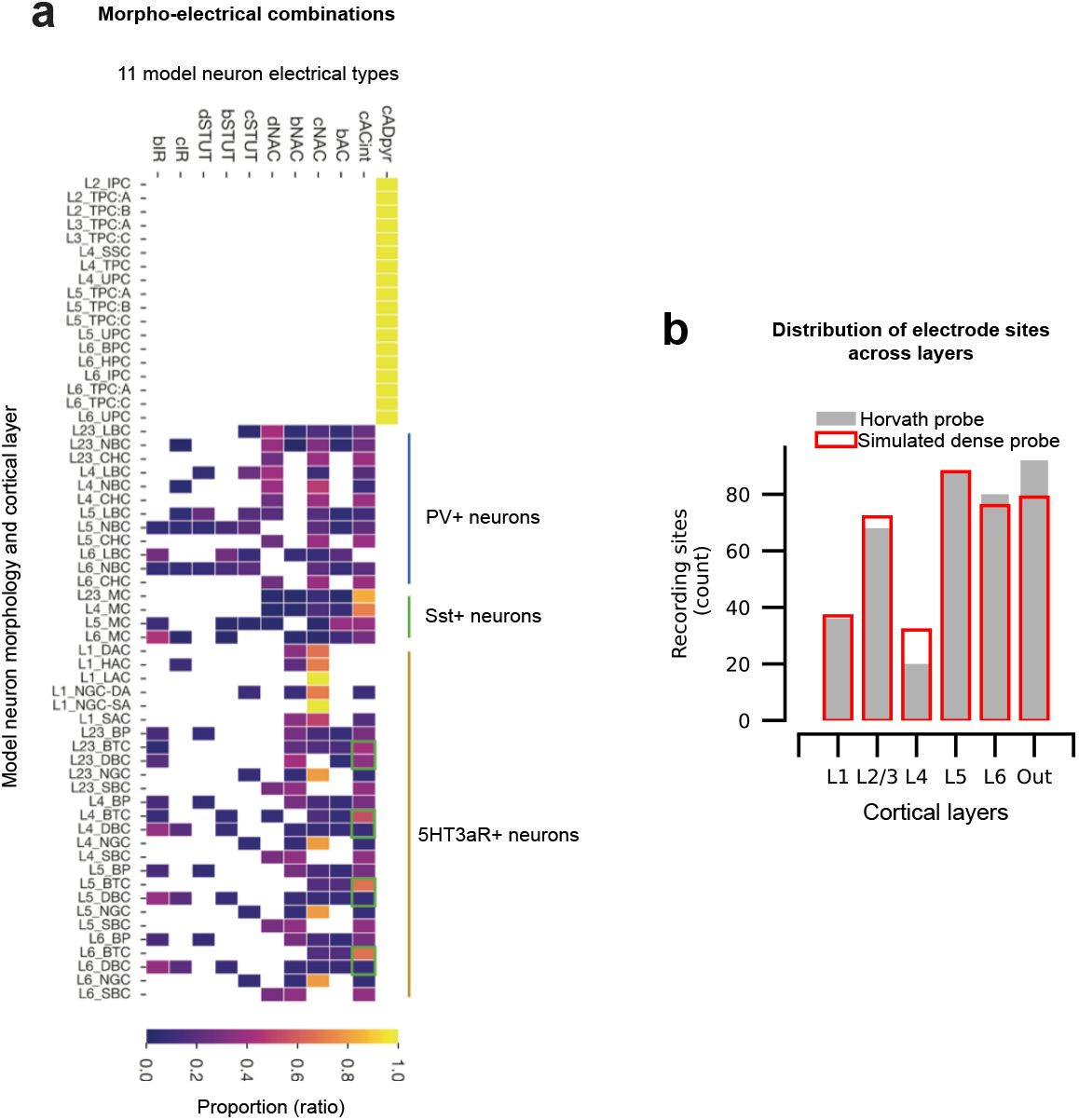
Model neuron combinations of morphological and electrical properties and coverage of the cortical column by the dense probe. **a**. morpho-electric types used in this study, i.e., the combinations of neuron morphologies, electrical firing patterns, layers and their proportions (heatmap colors). Inhibitory parvalbumin positive (PV+) cell class (blue bar), somatostatin positive cell class (Sst+, green) and 5HT3aR+ cell class (orange) are indicated by the vertical colored bars on the side. The cACint electrical type belongs to the Sst+ class (green boxes). The panel is adapted from [29]. **b**. The number of electrodes per layer of the Horvath setup [41] (filled grey bars) is plotted against the number of electrodes per layer of the simulated dense probe (empty red bars).

**Supplementary Fig. 2.**
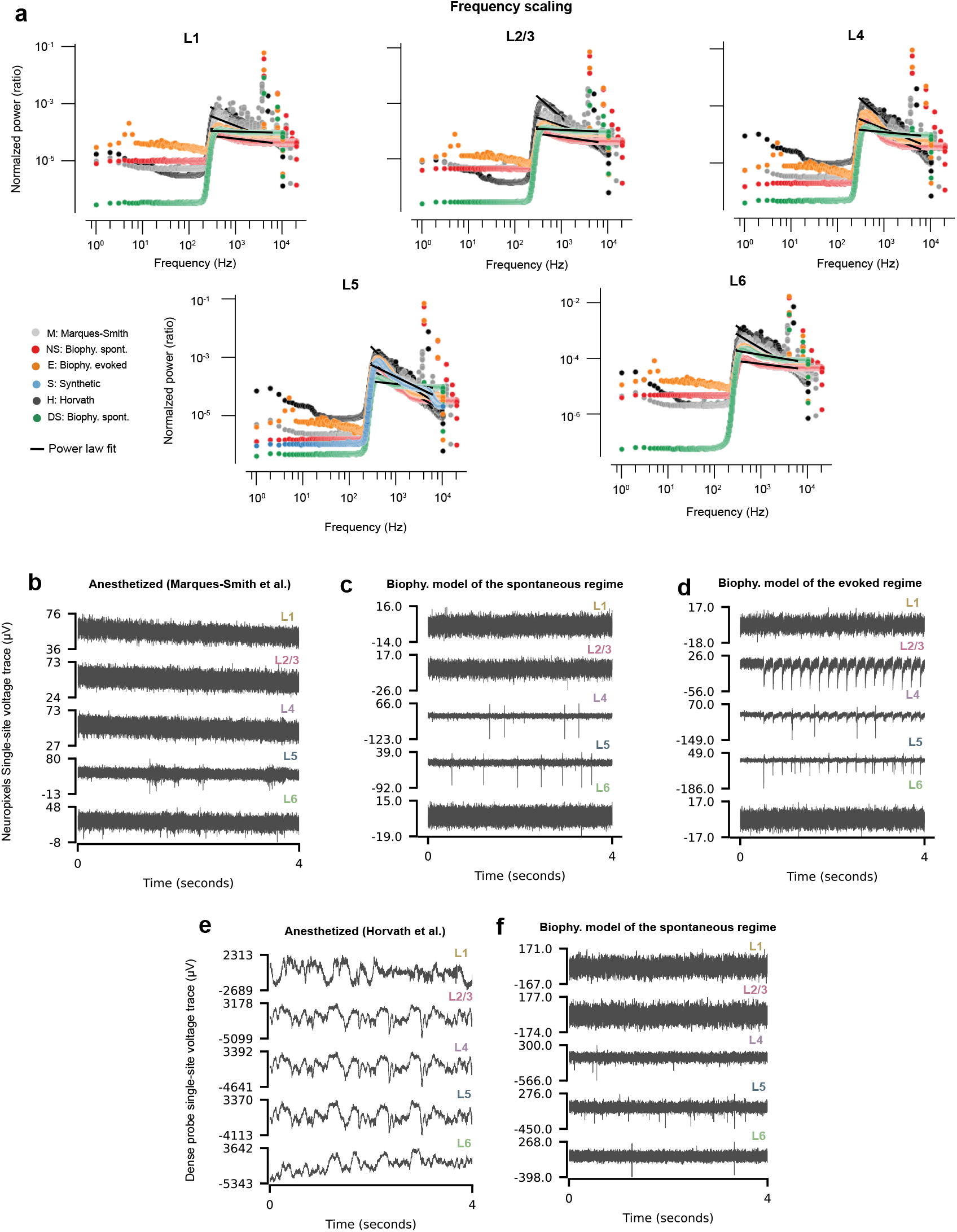
Validation of raw simulated traces. **a**. Median power spectral densities divided by the total power (PSDs) for high-pass filtered (with a cutoff at 300 Hz) and median subtracted traces are shown fitted with power-law functions (1*/f* ^*α*^) in the 0.3-6 KHz frequency band corresponding to spiking activity (black lines). Each panel displays the PSD of a cortical layer (L1: layer 1; L2/3: layer 2/3; L4: layer 4; L5: layer 5, L6: layer 6) and colors correspond to different experiments (see color legend) **b**. First 4 seconds of raw *in vivo* traces of Neuropixels from Marques-Smith data [8], **c**) biophysically simulated recordings in the **c**) spontaneous and **d**) evoked regime, **e**) Horvath dense probe recordings [41] and **f**) biophysically simulated recordings with Horvath dense probe, from the most superficial recording site of each cortical layers. Horvath traces displayed slow waves typically observed during slow wave sleep.

**Supplementary Fig. 3.**
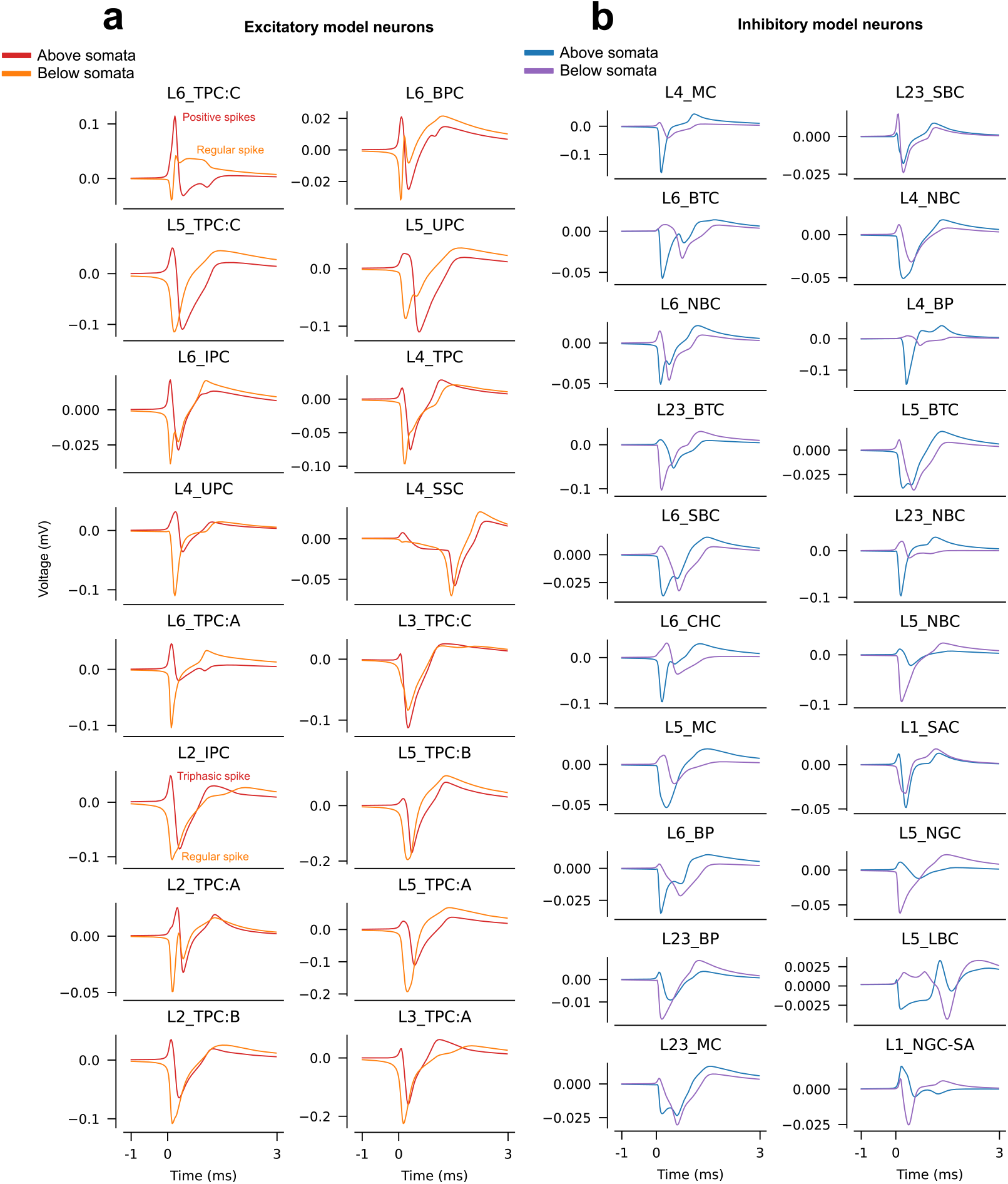
Changes in spike shapes with electrode location. Spike-triggered average of simulated recording traces from 16 excitatory **a**) and 20 inhibitory **b**) cells located within 50 µm of the Neuropixels probe and sorted by morphological type. The spike waveforms were recorded with two electrodes, one 20 µm above the cell somata (red in **a**, blue in **b**) and the other 20 µm below (orange in **a**, purple in **b**). For each morphological type, we display the cell with the highest firing rate.

**Supplementary Fig. 4.**
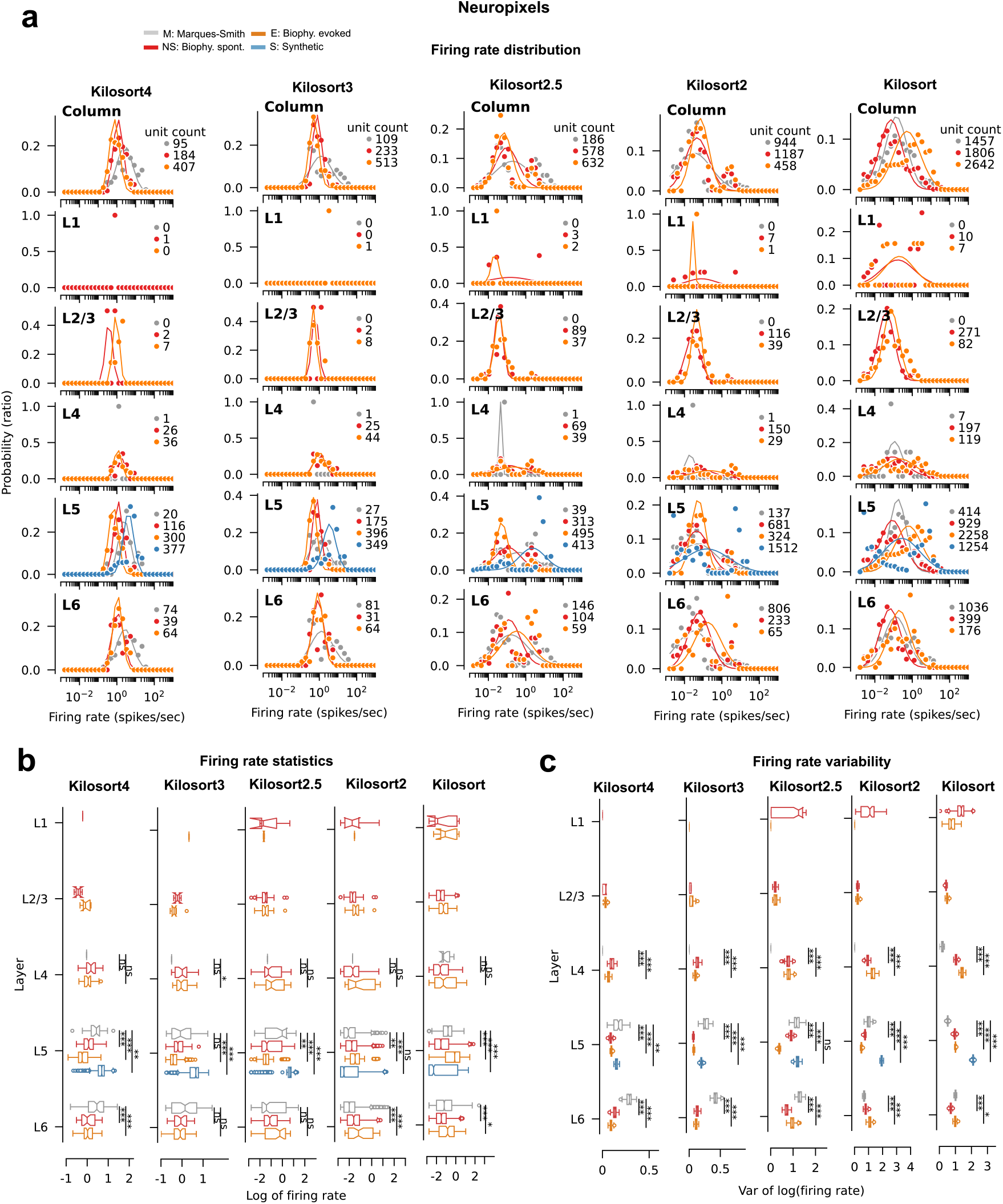
Validation of firing rate distributions. **a**. Distribution of firing rates of sorted single-units for each spike sorter (columns), layer (rows) and neuropixels experiment (colors). All units are reported for Kilosort, which does not classify single and ‘multi-unit’. The distributions were fitted with lognormal distributions (color-matched lines), which better matched the data for kilosort4 and 3 than for older sorters. **b**. Boxplots of log-transformed firing rates as in Fig. 4. Statistics are reported for the decimal logarithm of the firing rate. **c**.Boxplots of the variances of distributions of the firing rates generated from 100 bootstraps of the decimal logarithm of the firing rates. In **b** and **c**, boxplot notches represent the 95% confidence intervals. Statistical significance of the null hypothesis that the model sorted median firing rates and variances are the same as *in vivo* are reported with ns, *,**,***, which indicate *p >*= 0.05, *p <* 0.05, *p <* 0.01, *p <* 0.001 using the Mann-Whitney U test.

**Supplementary Fig. 5.**
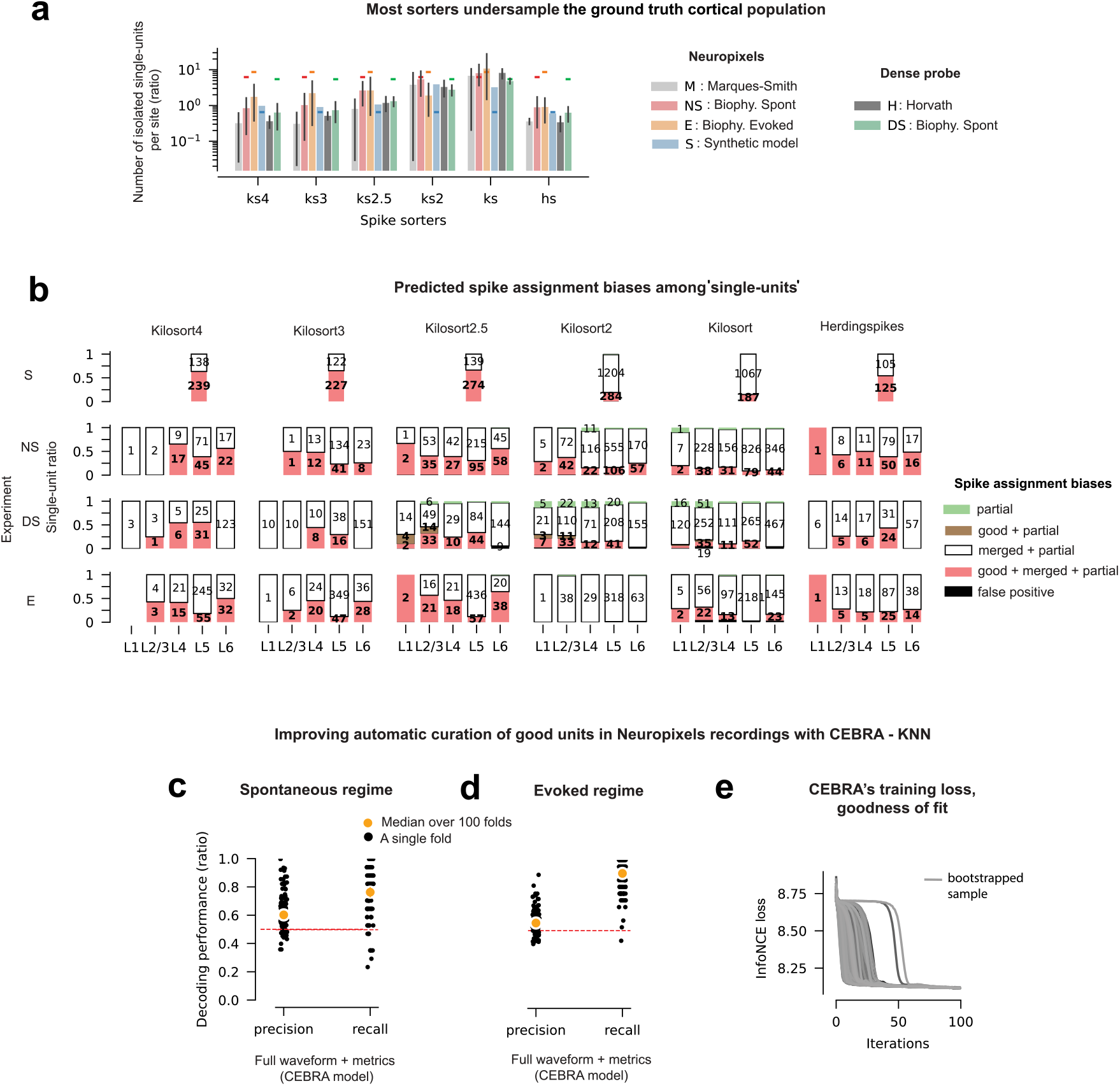
The model predicts significant undersampling and more diverse spike assignment biases for older spike sorters. **a**. ‘Single-unit’ yield per electrode across all spike sorters. The legend follows that in Fig. 5, with the median yield (bar) and 95% CI (error bars) shown across layers, and the theoretical yield averaged across layers indicated by the colored horizontal lines. **b**. Ratios of ‘single-unit’ spike assignment biases by layer (x-axis), spike sorter (column panels), and ground truth simulation (row panels). The overlaid numbers represent the single-unit counts. Also shown are the precision and recall of a model incorporating both the full spike waveforms and the ten spiking quality metrics from Fig. 5g. The model uses CEBRA, an autoencoder that reduces the dimensionality of the full waveforms to three dimensions, followed by a k-nearest neighbor classifier to distinguish ‘good’ and degraded units based on the embeddings and quality metrics. Performance is reported for the **c**) spontaneous and **d**) evoked biophysical simulations of Neuropixels recordings. **e**. InfoNCE loss functions of the model for all bootstrapped samples. All cross-validated folds of CEBRA converged within 60 iterations, indicating sufficient training.

**Supplementary Fig. 6.**
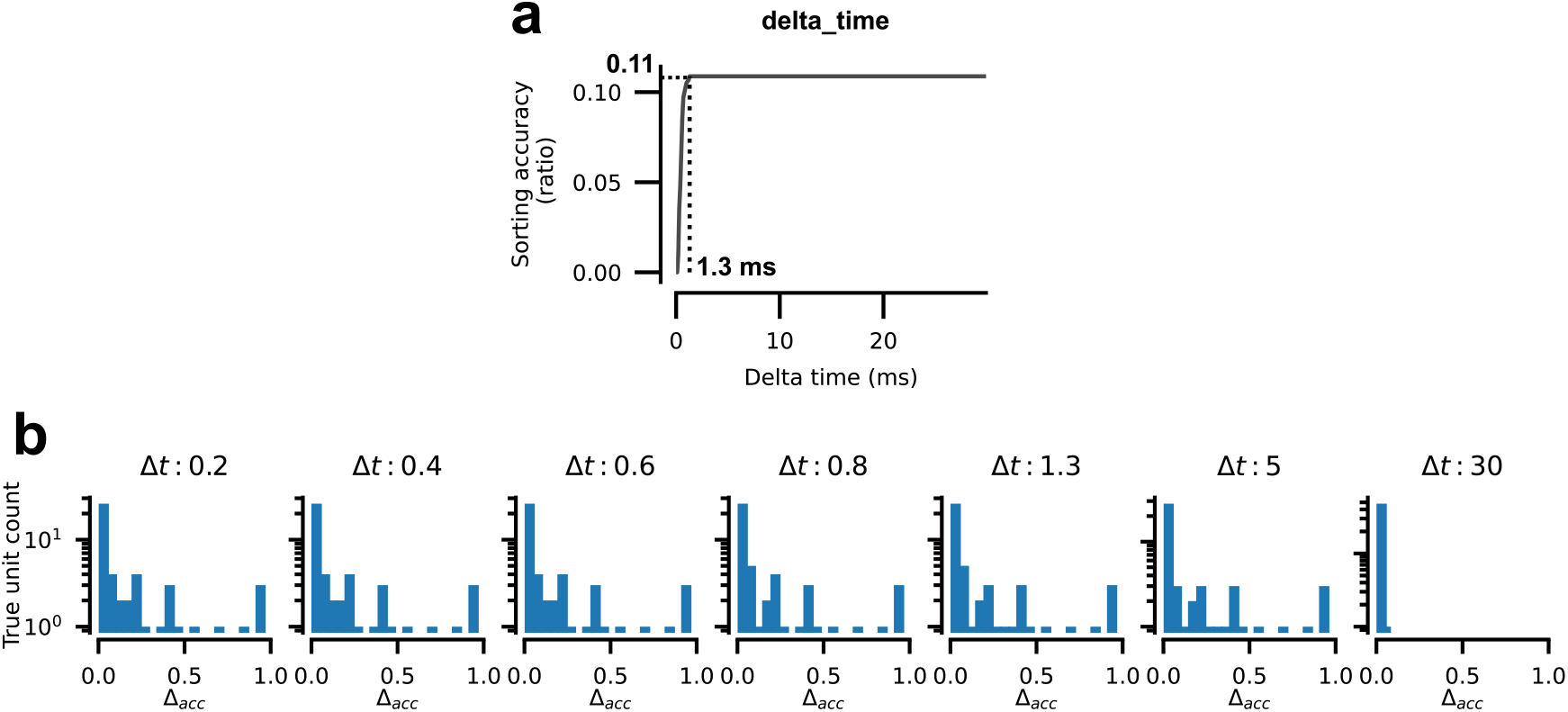
Sorting accuracy. **a**. Effect of the delta time (Δ*t*) parameter on sorting accuracy, for 10 min recording of the simulation of Neuropixels recordings sorted with Kilosort4. Δ*t* is the time window around each ground truth spike within which coincidences were detected between sorted and ground truth spikes. Sorting accuracy is maximized for a Δ*t* duration of 1.3 ms. **b**. Distribution of ground truth neuron sorting accuracies for zero shift versus long constant shifts of ground truth spike timestamps. When Δ*t* becomes too long (e.g., Δ*t* = 30 ms), the accuracy distribution shows a single peak near 0, indicating chance-level performance. For meaningful Δ*t* (from 0.2 to 5 ms), the distribution displays more than one peak: peaks for correctly detected units and a peak near chance level for missed units. A Δ*t* of 1.3 ms is not overly lenient: it does not produce a single peak near 0, indicating that high scores are assigned for random matches, but produces several high score peaks.

**Supplementary Fig. 7.**
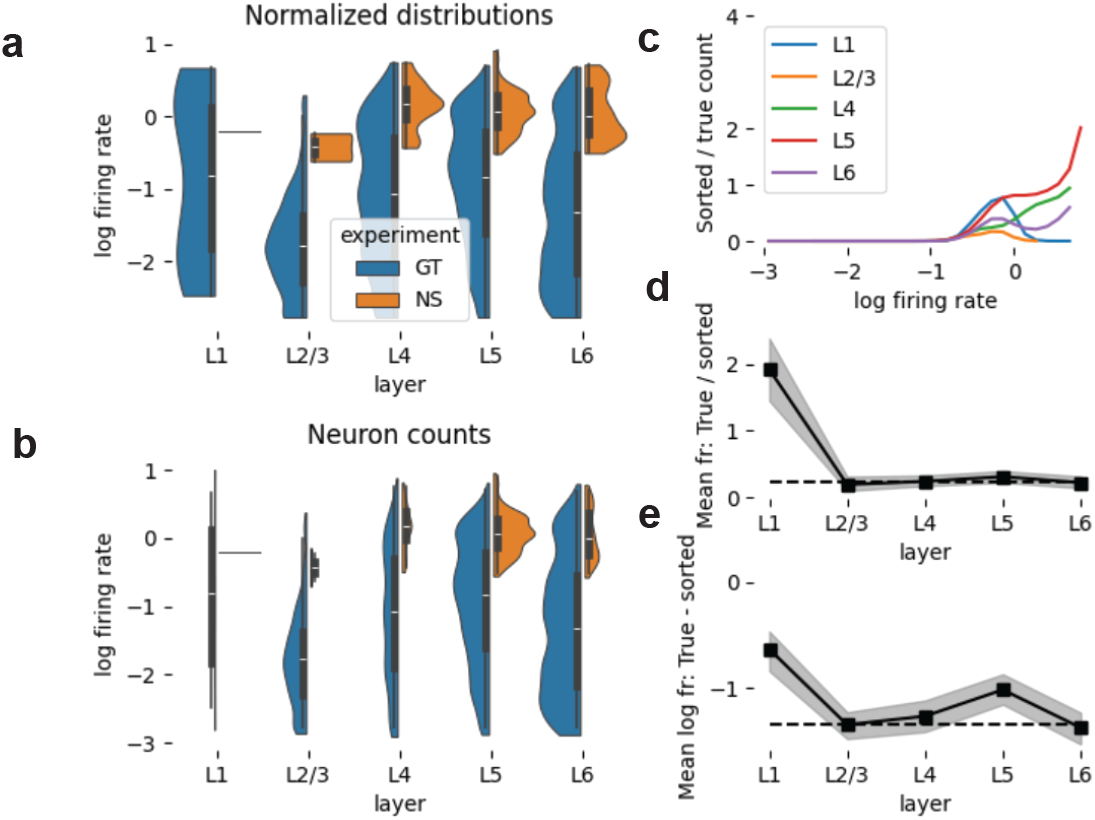
Ground truth firing rate distribution. **a**. Violinplots of the distributions of sorted (orange) and true (blue) firing rates in individual layers for the spontaneous biophysical simulations of Neuropixels recording. True firing rates are based on all neurons within 50*µm* of the recording electrode. Widths of the violins normalized for each class. **b**. as **a**, but with non-normalized widths, i.e. the widths are directly proportional to the number of neurons in each bin. **c**. Ratio of sorted divided by true neuron count in each logarithmic firing rate bin. Colors indicate results for individual layers. **d**. Ratio of true divided by sorted mean firing rates in each layer. Grey area indicates a 90% confidence interval based on bootstrapping. **e**. as D, but instead the difference between the logarithm of the means is indicated.

**Supplementary Fig. 8.**
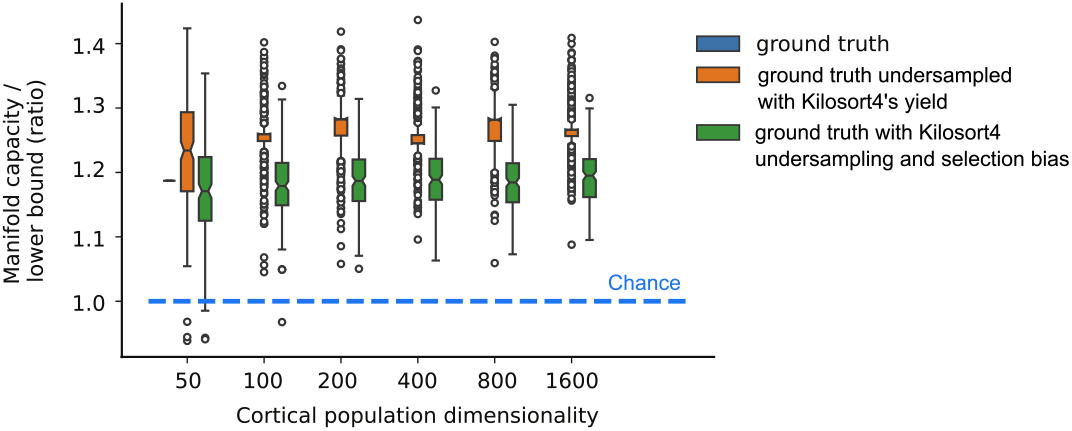
Information capacity remains stable despite changes in neuronal feature dimensionality. The normalized information capacity of randomly undersampled ground truth neurons—either matched to Kilosort4’s yield or sampled to reproduce Kilosort4’s unit-type distribution is plotted against increasing dimensionality of a Gaussian random projection.

